# Tracing the Ancestry of Operons in Bacteria

**DOI:** 10.1101/212886

**Authors:** Huy Nguyen, Ashish Jain, Oliver Eulenstein, Iddo Friedberg

## Abstract

**Motivation:** Complexity is a fundamental attribute of life. Complex systems are made of parts that together perform functions that a single component, or subsets containing individual components, cannot. Examples of complex molecular systems include protein structures such as the *F*_1_*F*_*o*_-ATPase, the ribosome, or the flagellar motor: each one of these structures requires most or all of its components to function properly. Given the ubiquity of complex systems in the biosphere, understanding the evolution of complexity is central to biology. At the molecular level, operons are a classic example of a complex system. An operon’s genes are co-transcribed under the control of a single promoter to a polycistronic mRNA molecule, and the operon’s gene products often form molecular complexes or metabolic pathways. With the large number of complete bacterial genomes available, we now have the opportunity to explore the evolution of these complex entities, by identifying possible intermediate states of operons.

**Results:** In this work, we developed a maximum parsimony algorithm to reconstruct ancestral operon states, and show a simple vertical evolution model of how operons may evolve from the individual component genes. We describe several ancestral states that are plausible functional intermediate forms leading to the full operon. We also offer **R**econstruction **o**f **A**ncestral **G**ene blocks **U**sing **E**vents or ROAGUE as a software tool for those interested in exploring gene block and operon evolution.

**Availability:** The software accompanying this paper is available under GPLv3 license in:

https://github.com/nguyenngochuy91/Ancestral-Blocks-Reconstruction.

All figures in this paper are available in enlarged downloadable form from:

https://github.com/nguyenngochuy91/Ancestral-Blocks-Reconstruction/tree/master/images

## Introduction

The evolution of complex systems is an open problem in biology (Wagner and Altenberg, 1996; Bonner, 1988; Pál and Papp, 2017), and has recently been studied intensively in genomes (Adami, Ofria, and Collier, 2000; Lynch and Conery, 2003; Koonin and Dolja, 2006). To better understand how complex systems evolve, we focus on the problem of the evolution of orthologous gene blocks and operons in bacteria. Orthologous gene blocks or *orthoblocks* are sequences of genes co-located on the chromosomes of several species, whose evolutionary conservation is apparent. Operons can be viewed as a special case of gene blocks where the genes are co-transcribed to polycistronic mRNA and are often associated with a coherent function, such as a metabolic pathway or a protein complex. Several models have been proposed to explain gene block and operon evolution. It may very well be that the models are not mutually exclusive, and different operons may evolve by different models, or indeed a single operon may be the result of the combination of several models (Horowitz, 1945; Stahl and Murray, 1966; Omelchenko *et al.*, 2003; Lawrence and Roth, 1996; Fani, Brilli, and Li ò, 2005; Price, Arkin, and Alm, 2006; Alm, Huang, and Arkin, 2006; Koonin, 2009; Hsiao *et al.*, 2005; Goldberg, Rost, and Bromberg, 2016; Bush *et al.*, 2018).

Previously, we proposed a method that explains the evolution of orthoblocks and operons as a combination of events that take place in vertical evolution from common ancestors (Ream, Bankapur, and Friedberg, 2015). In the evolution of an orthoblock, the different gene blocks may gain or lose genes, have genes duplicated, or have them split off (Figure 1 and Table 1). By determining the frequency of the events for any orthoblock in a studied clade, we can determine a cost for each event, and thus create a cost function to determine an optimal vertical path for the evolution of orthoblocks. We used the cost function to determine the conservation of some operons and orthoblocks in proteobacteria, and show that orthoblocks that perform cellular information processing (such as mRNA translation) are more conserved than those that are associated with adaptation to specific environments.

**Table 1.** All pairwise events for the orthoblocks shown in Figure 1.

|  | A | B | C | D | E |
| --- | --- | --- | --- | --- | --- |
| A |  |  |  |  |  |
| B | split, deletion |  |  |  |  |
| C | deletion | split |  |  |  |
| D | duplication, deletion, split | duplication, 2× split, | duplication, split |  |  |
| E | duplication, deletion | duplication, split | duplication | split |  |

**Fig 1.**
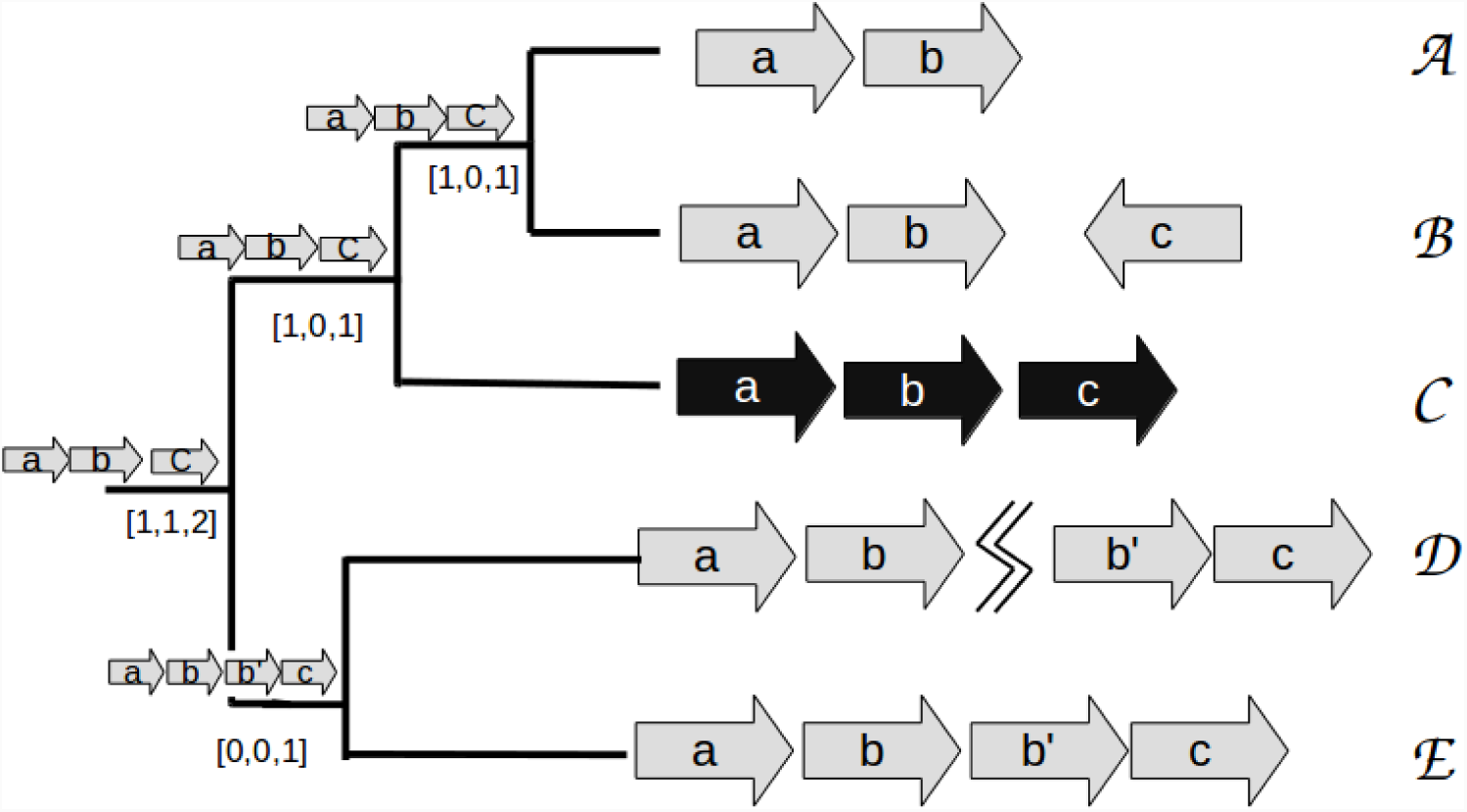
Orthoblocks from species A-E are arranged in a species phylogenetic tree. Species C has an experimentally-determined operon (Black arrows), and serves as the reference taxon. The orthologs in species A,B,D, and E were determined as explained the in the text. The events between C and all other species for this orthoblock are: A-C: deletion (of gene c) B-C: split (of gene c) C-D: duplication (of b) and split (jagged line) C-E: duplication (of b) The full list of the pairwise events between all species is in Table 1. The tree’s inner nodes show proposed intermediate states in the operon’s evolution. The numbers in the brackets are a 3-tuple showing the cumulative count of events going from the leaf nodes to the tree root: [deletions, duplications, splits]. The way these ancestral states are determined is elaborated below

In this study we use the orthoblock evolution distance function to reconstruct ancestral gene blocks. Reconstructing plausible ancestral states of extant gene blocks and operons can help us understand how they evolve, identify possible functional intermediate states, and determine which forces might affect their evolution. The rest of this paper is structured as follows: first, we describe our approach, elaborating on the two algorithms we developed. We then present and discuss the results using our algorithms to reconstruct the ancestral states of orthoblocks in a clade of Gram-negative bacteria and a clade of Gram-positive bacteria. This reconstruction involves orthoblocks comprising genes orthologous to those found in operons in *Escherichia coli* and in *Bacillus subtilis*, respectively. Our reconstructions of ancestral states show that: (1) some operons can rapidly evolve independently in several branches in their respective clades, suggesting that positive selection plays a major role in the evolution of gene blocks in bacteria; (2) other operons are highly conserved, their evolution predating the last common ancestor of the clades we chose, (3) some ancestral state can plausibly be described as intermediate functional forms, and (4) some operon conservation is sporadic and cannot be explained solely by vertical transmission suggesting horizontal gene transfer.

## Materials and Methods

### Definitions

#### Gene block-based evolutionary events, and event-based distances

A **reference taxon** is a taxon where operons have been identified by experimental means. Here we use*E. coli* K-12 MG1655 (NC 000913) and *B. subtilis* str. 168 (NC 000964) as reference taxa. The reference taxon serves as a standard of truth to determine if the genes on a suspected orthoblock do indeed reside, at least in one species, in an operon or a similar co-regulated gene block. **Neighboring genes**: two genes are considered neighboring if they are ≤ 500 bp apart and on the same strand, variations in threshold between 300 and 700 bp showed little difference in previous studies by us (Ream, Bankapur, and Friedberg, 2015). A **gene block** comprises no less than two neighboring open reading frames (ORFs). **Orthoblocks**, gene blocks that are orthologous, are defined as follows: two organisms have orthoblocks when each organism must have at least two neighboring genes that are homologous to genes in a gene block in the reference taxon’s genome. An **event** is a change in the gene block between any two species with homologous gene blocks.

We identify three types of pairwise events between orthoblocks in different taxa: splits, deletions, and duplications. The *event-based distance* between any two orthoblocks is the sum of the minimized count of splits, duplications, and deletions, which is elaborated upon in **Orthoblock Distance Functions**. See Figure 1. The terms reference taxa, neighboring genes, gene blocks, events, and orthoblocks are elaborated upon in (Ream, Bankapur, and Friedberg, 2015).

#### Choosing species

The species tree for each clade was built using *rpoB* as the species marker. For the study of Gram negatives with *E. coli* as a reference species, we use the group of taxa from (Fani, Brilli, and Lio, 2005). For the study of Gram positives with *B. subtilis* as the reference species, we use the Phylogenetic Diversity Analysis program (PDA)(Chernomor *et al.*, 2015; Faith, 1992) to select 33 equidistant species. Note that other species markers can be used, and the choice for those may vary depending on the number of species analyzed and the phylogenetic distances. While in this study we used *rpoB*, ROAGUE can use the input from any species tree provided.

#### Orthoblocks in Phylogenetic Trees

For each orthoblock studied, we use a phylogenetic species tree *T* comprising a set of extant species related to a reference taxon. Each leaf node *ν* in *T* contains the orthologs to the genes in an operon in the reference species. For any two genes *a* and *b*, if the chromosomal distance is less than 500 bp, the genes will be written as *ab*. If the distance is greater than 500 bp, they are written with the separator character ‘|’ thus: *a*|*b*. For a species tree *T,* we define *V* (*T*), *E*(*T*), *L*(*T*) as the set of nodes, edges, and leaves of *T* respectively. In addition, we denote *T*_*ν*_ to be a subtree of *T* rooted at ν ∈ *V* (*T*). *T*_*ν*_ is obtained by pruning *T*_*ν*_ from *T*.

Given a reference operon ℴ, we define 𝒢 := {*x*_1_, *x*_2_, *x*_3_, *…, x*_*n*_} be the set of gene of ℴ. We denote a gene block ℬ over 𝒢 is a non empty multiset of 𝒢, 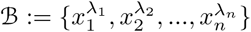 where *x*_*i*_ ∈ 𝒢, λ_*i*_ ∈ ℕ. We define the set of genes in gene block ℬ as Gene(ℬ) := {*x*_*i*_ λ_*i*_ ≥ 1}. We also define duplication gene set of a gene block ℬ as Dup(ℬ) := {*x*_*i*_ λ_*i*_ ≥ 2}. An orthoblock *O* is a set of blocks that either empty or contains at least one gene block of size at least 2. Given a gene block ℬ and a gene set *G* over 𝒢, we define 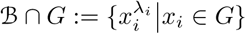

Given a species tree *T* and a reference operon ℴ, for node *ν* ∈ *V* (*T*), let *O* be the orthoblock assigned to *ν*, we define:

1. *I*_*g*_(*ν*): the identity function of gene *g* in *O*.
2. *ν.gene*[*g*]: the set that represents whether to include gene *g* in *O*. There are only 3 possible cases.
  a. *ν.gene*[*g*] = {1} : this means that gene *g* has to be in *O*.
  b. *ν.gene*[*g*] = {0} : this means that gene *g* can not be in *O*.
  c. *ν.gene*[*g*] = {0, 1} : this means that gene *g* can either be in *O* or not in *ν*.
3. *ν.dup*[*g*]: the set that represents the duplication status of gene *g* in *O*. There are only 3 possible cases.
  a. *ν.dup*[*g*] = {1} : this means that gene *g* has to be duplicated in *O*.
  b. *ν.dup*[*g*] = {0} : this means that gene *g* can not be duplicated in *O*.
  c. *ν.dup*[*g*] = {0, 1} : this means that gene *g* can either be duplicated or not in *O*.
4. *Gene*(*O*): the set of genes of *O*. *Gene*(*O*) := ∪_ℬ∈*O*_ Gene(ℬ)
5. *Dup*(*O*): the set of gene that is duplicated in some gene blocks of *O*. *Dup*(*O*) := ∪_ℬ∈*O*_ Dup(ℬ)
6. *FREQ*_*g*_(*ν*): The proportion of *L*(*T*_*ν*_) that contains gene *g*.
7. *DUP*_*g*_(*ν*): The proportion of *L*(*T*_*ν*_) that contains a duplication of gene *g*.

### Orthoblock Distance Functions

The distances between any two orthoblocks *O, O’* are defined as follows:

1. *Split distance* (*d*_*s*_) is the absolute difference in the number of relevant gene blocks between the two taxa. We define *Rel*(*O, O’*) is the set of gene block from *O* where each gene in each gene block has to appear in *O’* at least once. Formally, *Rel*(*O, O’*) := ⋃_𝔅∈*O*_ (𝔅 ∩ *Gene*(*O’*)). The split distance can be formalized as:

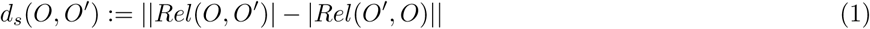

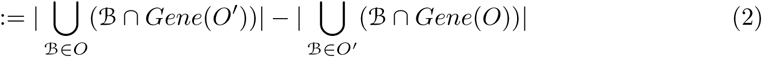 Example: for the reference gene block with genes (abcdefg), genome A has blocks *O* := ((*ab*), (*def*)) and genome B has *O’*:= ((*abc*), (*de*), (*fg*)). We then compute the relevant gene blocks *Rel*(*O, O’*) = ((*ab*), (*def*)) and *Rel*(*O’, O*) = ((*ab*), (*de*), (*f*)) (removing genes *c, g*). Therefore, *d*_*s*_(*O, O’*) = |2 *-* 3| = 1.
2. *Duplication distance (d*_*u*_) is the pairwise count of duplications between two gene blocks. We define *Dif* (*O, O’*) as the set of duplicated genes of gene block *O*, so that these genes also appear in *O’* but are not duplicated in *O’*. Formally, *Dif* (*O, O’*) := (*Dup*(*O*) ∩ *Gene*(*O’*)) *\ Dup*(*O’*). Here, our gene blocks are guarantee to have at most one duplication of each gene for each block. We formalize the duplication distance as:

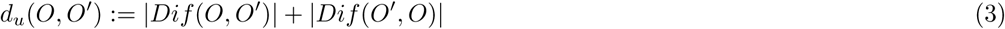

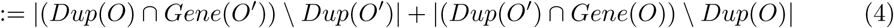 Example: For a reference gene block (*abcde*), genome A has gene block *O* = ((*abd*)) and genome B has gene block *O’* = ((*abbcc*)), respectively. The ortholog of gene *O*_*b*_ is duplicated in genome B, creating a duplication distance *d*_*u*_(*O, O’*) of 1. However, since gene *c* does not exist in *O*, it has no bearing on the duplication distance between the homologous gene blocks *O* and *O’*. We then compute *Dif* (*O, O’*) = ∅ and *Dif* (*O’, O*) = {*b*}. Therefore, *d*_*u*_(*O, O’*) = 0 + 1 = 1.
3. *Deletion distance (d*_*d*_) is the difference in the number of orthologs that are in the homologous gene blocks of the genome of one organism, or the other, but not in both. In short, it is the symmetric difference between the set of orthologous genes of the two gene blocks *O, O’*. We formalize the deletion distance as:

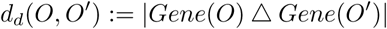 In addition, the deletion distance can also be defined using the identity function:

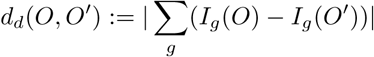 Example: For a reference gene block (*abcde*), genome A has gene block *O* = ((*abd*)) and genome B has gene block *O’* = ((*abce*)), respectively. Since there are only genes *a, b* that appear in both genomes, *d*_*d*_(*O, O’*) = |{*a, b, d*} Δ {*a, b, c, e*}| = |{*d*}| + |{*c, e*}| = 3

Intuitively, the duplication distance, split distance, and deletion distance are not interdependent. However, as each distance contains the variables *Gene*(*O*), *Gene*(*O’*), the three distances are actually interdependent. Using the three distance functions above, we define the total distance between any two homologous gene blocks *O, O’* as:

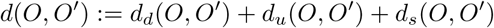

### Problem Definition

Let *T* be a tree, and *G* be the set of genes in a reference operon. We define Ω as the set of all possible orthoblocks over gene set *G*. Let λ : *L*(*T*) ↦ Ω be the labeling of *L*(*T*) (assign orthoblocks from Ω to the leaf nodes of *T*, this can include empty orthoblocks). We define the function 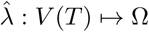 to be an extension of λ on *T* if it coincides with λ on the leaves of *T* (assign an orthoblock to each node of *T*). If 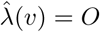, we say that vertex *v* is labelled with orthoblock *O*. Given a labelling 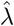 and an edge (*u, v*) ∈ *E*, we define the distance between the two labellings of the endpoints *u, v* as 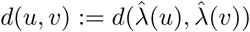 and the total distance function as 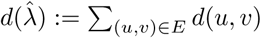.

The Maximum Parsimony problem is now defined as follows: given a tree *T*, an operon gene set *G*, the orthoblock set Ω and a leaf labeling *λ*, find a labeling 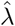 that minimizes 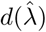

Here we explore two related Maximum Parsimony heuristic approaches, local and global, to reconstruct ancestral gene blocks.

### Local Maximum Parsimony

Because the three distance measures are interdependent, the local parsimony problem is not trivial. In the following example, we demonstrate why it is difficult to infer a parent from children in the most parsimonious way.

Given an inner node *v* and its two child nodes *v*_1_ and *v*_2_, let *O* be the gene block to be assigned to *v*. Consider the orthoblocks *O*_1_ and *O*_2_ of *v*_1_ and *v*_2_ respectively as:

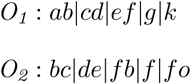

We define the set of genes that appear in both *O*_1_ and *O*_2_ as *S* = {*b, c, d, e, f*}, and the union gene set of *O*_1_ and *O*_2_ as *G* = {*a, b, c, d, e, f, g, k, o*}. Any gene *i* ∈ *S* will contribute a deletion distance of 2 to *d*_*d*_(*O, O*_1_) + *d*_*d*_(*O, O*_2_) if *O* does not contain gene *i*. Any gene *i* ∈ *G* but *i* ∉ *S* will contribute a deletion distance of 1 to *d*_*d*_(*O, O*_1_) + *d*_*d*_(*O, O*_2_) if *O* either has it or not. Hence, including all genes from *S* in *O* gives us deletion distance: *d*_*d*_(*O, O*_1_) + *d*_*d*_(*O, O*_2_) = 4, which is the minimum deletion distance. On the other hand, if we just want to minimize the split distance, the most naive way is not to include any genes in *O*. Then, *Rel*(*O, O*_1_) = *Rel*(*O, O*_2_) = ∅, therefore *d*_*s*_(*O, O*_1_) + *d*_*s*_(*O, O*_2_) = 0. However, if we choose to do it this way, our deletion distance becomes large (*d*_*d*_(*O, O*_1_) + *d*_*d*_(*O, O*_2_) = 10). Apparently, decreasing split distance might increase deletion distance and vice versa.

If we focus on minimizing the deletion distance, then *Gene*(*O*) = *S*, which means that *O* has to include all genes in *S*. Then, the relevant gene blocks of *O*_1_, *O*_2_ to *O* respectively become:

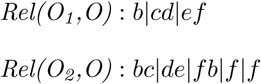

The split distance of *O*_1_, *O*_2_ is *d*_*s*_(*O*_1_, *O*_2_) = |5 *-* 3| = 2. If we remove gene *f* from *Gene(O)*, the relevant gene blocks of the two children to *u* become:

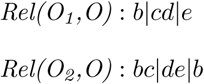

Hence, by setting our gene block *O* as either *Rel*(*O*_1_, *O*) or *Rel*(*O*_2_, *O*), the deletion distance increased by 2 because we excluded gene *f* which is in *S*; however, the split distance also decreased by 2. Therefore, the new deletion distance is *d*_*d*_(*O, O*_1_) + *d*_*d*_(*O, O*_2_) = 6, and the new split distance is *d*_*s*_(*O, O*_1_) + *d*_*s*_(*O, O*_2_) = 0.

Consider another possibility: if we include gene *g* in *Gene*(*O*), this will not increase the deletion distance. The relevant gene blocks of the two children to *u* become:

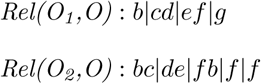

By setting *O* := *b*|*cd*|*ef* |*g*, the new split distance is *d*_*s*_(*O, O*_1_) + *d*_*s*_(*O, O*_2_) = 0 + 1 = 1 and the deletion distance is *d*_*d*_(*O, O*_1_) + *d*_*d*_(*O, O*_2_) = 4. Therefore, we achieve a lower aggregate sum of deletion and split distances (5 compared to 6). We can keep on adding, or removing genes that only appear in one taxon. This process requires iterations through all the subsets of the symmetrical difference set *Gene*(*O*_1_)Δ*Gene*(*O*_2_), which will take exponential time. We therefore provide a heuristic approach that guarantees minimum deletion and duplication distances, but not split distances.

Briefly, the local approach focuses on finding the optimal parent ancestral gene block given its child gene blocks. For each internal node *u*, let *u*_1_ and *u*_2_ be its two children. The intuition is that we have to include a gene in the parent if both of the children have it. However, greedily propagating an included gene up a tree may cause predicting its ancestral existence into deeper internal node than is warranted. To check this problem, at each tree vertex *v*, for each gene *g*, we introduce a correction by checking the fraction of the leaf nodes that contain *g*. Since gene loss tends to happen more often than gene gain, we use a threshold of 0.5 to indicate whether *v* contains gene *g* or not. We present a greedy local optimization algorithm as follows. See Figure 2 for a visualization of the process.

**Fig 2.**
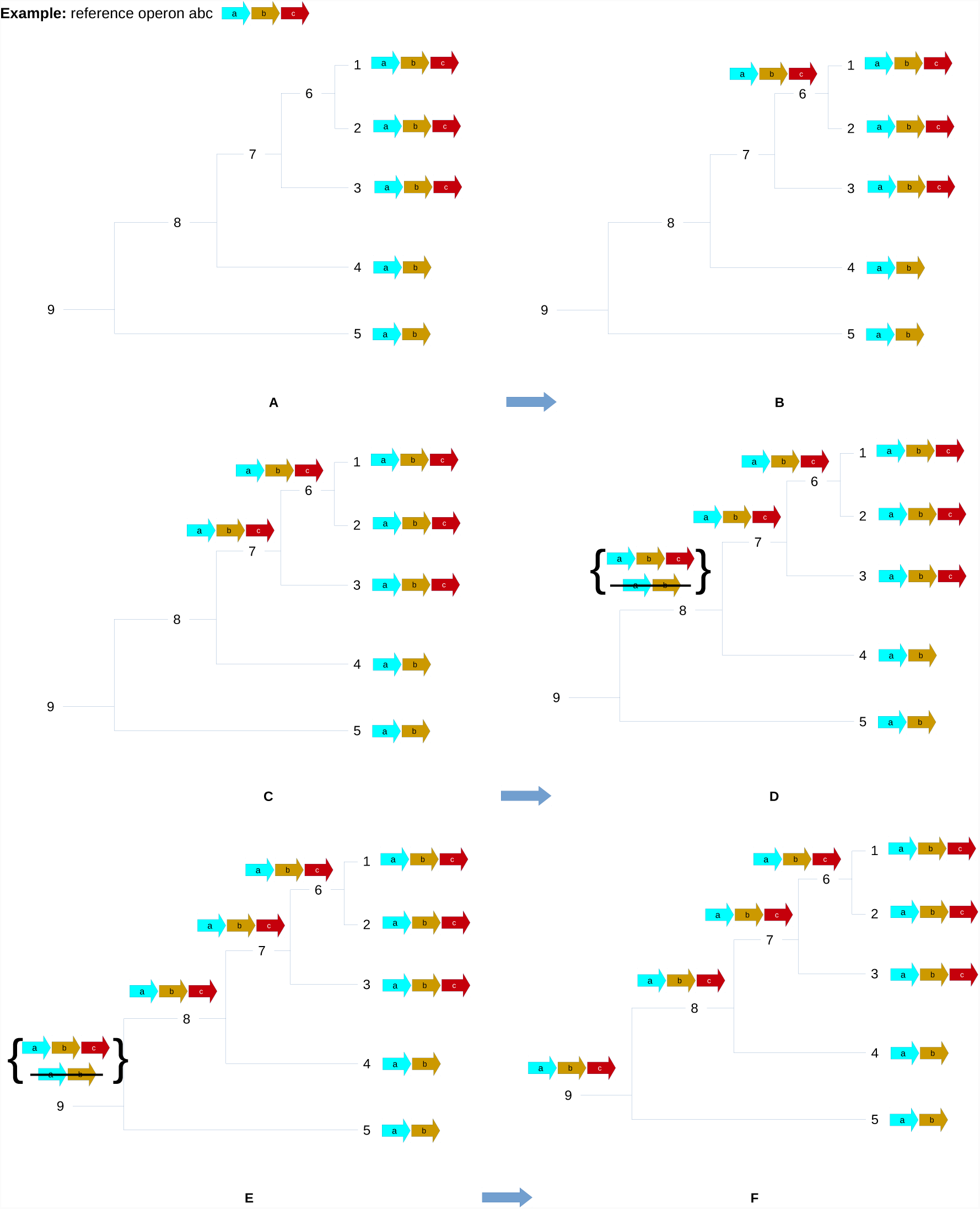
(Previous page.) A simplified example of ancestral reconstruction using local maximum parsimony. Consider a tree with structure as in panels A. In each panel, **1,2,3,4,5** are the extant nodes that are assigned with gene blocks **abc, abc, abc, ab, ab** respectively, and **6,7,8,9** are the inner nodes. The local algorithm traverses the tree bottom-up. In panel A,B,C, the gene block reconstruction of node **6,7** is **abc** (**1,2,3** all have gene block **abc**). In panel D, node **8**, there are 2 best candidates for the gene block reconstruction. However, we chose to include gene **c** since *FREQ*_*c*_(**8**) = 3*/*4 = .75 *> .*5. Hence, node **4** is assigned with gene block **abc**. In panel E, at node **9**, there are three best candidates. We chose to assign gene block **abc**. The reason is that *FREQ*_*c*_(**5**) = *FREQ*_*c*_(**5**) = 3*/*5 = .6 *> .*5.

#### Algorithm 1 Local cost function minimization for reconstructing ancestral nodes

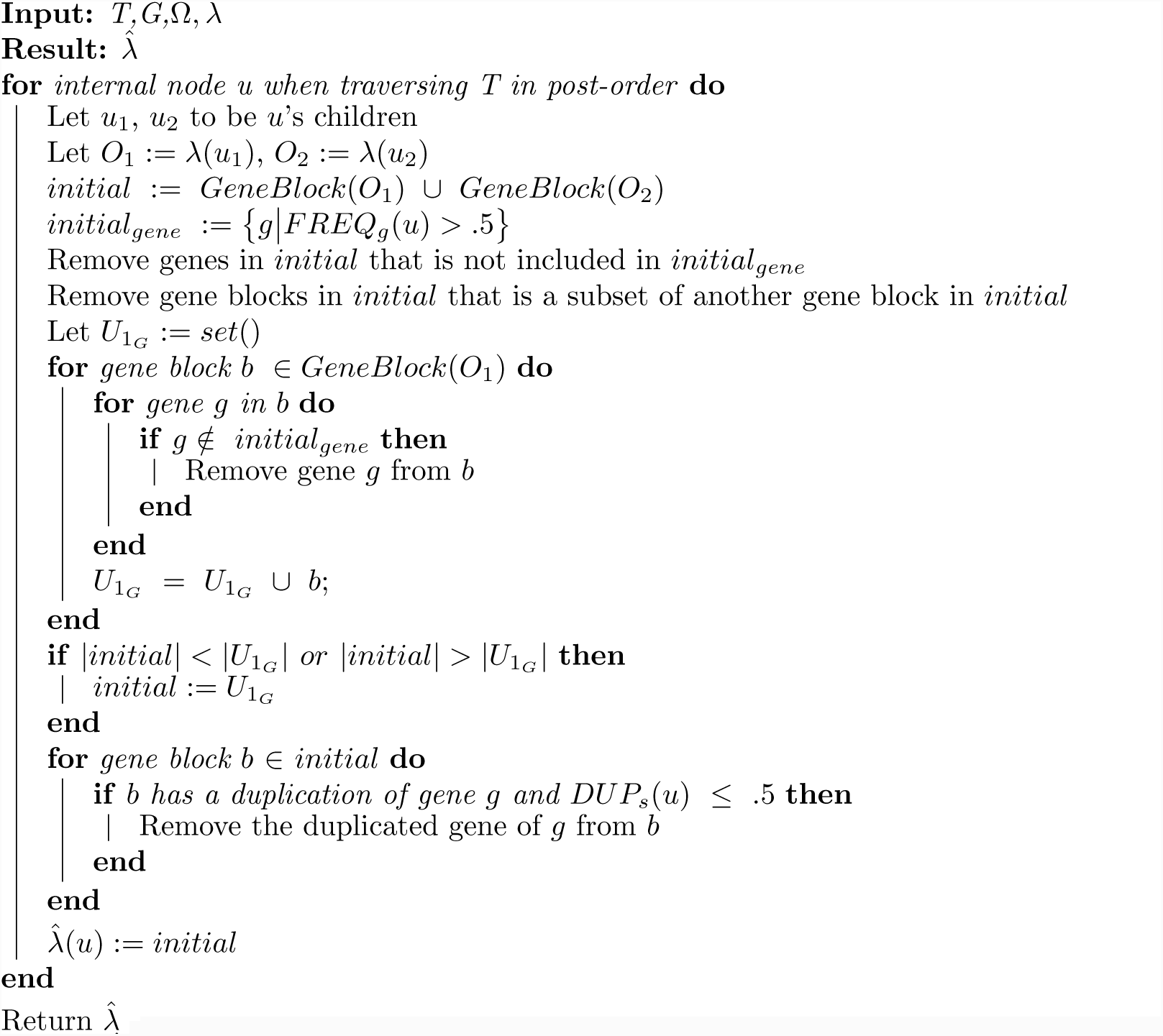

### Global Maximum Parsimony

Here, we try to achieve minimal global deletion and duplication distances. Intuitively, for each node *v* and for each gene *g* in the reference operon, we decide whether gene *g* would appear in the orthoblock we will assign to *v*. To do this, we use dynamic programming. By traversing the phylogenetic tree bottom-up and top-down, we determine the occurrence of each gene in the reference operon for each node *v*. We also determine whether a gene should be duplicated in the same manner. In term of split distances, we generate the relevant gene blocks of the two children given the set of gene to be included. See also Figure 3.

**Fig 3.**
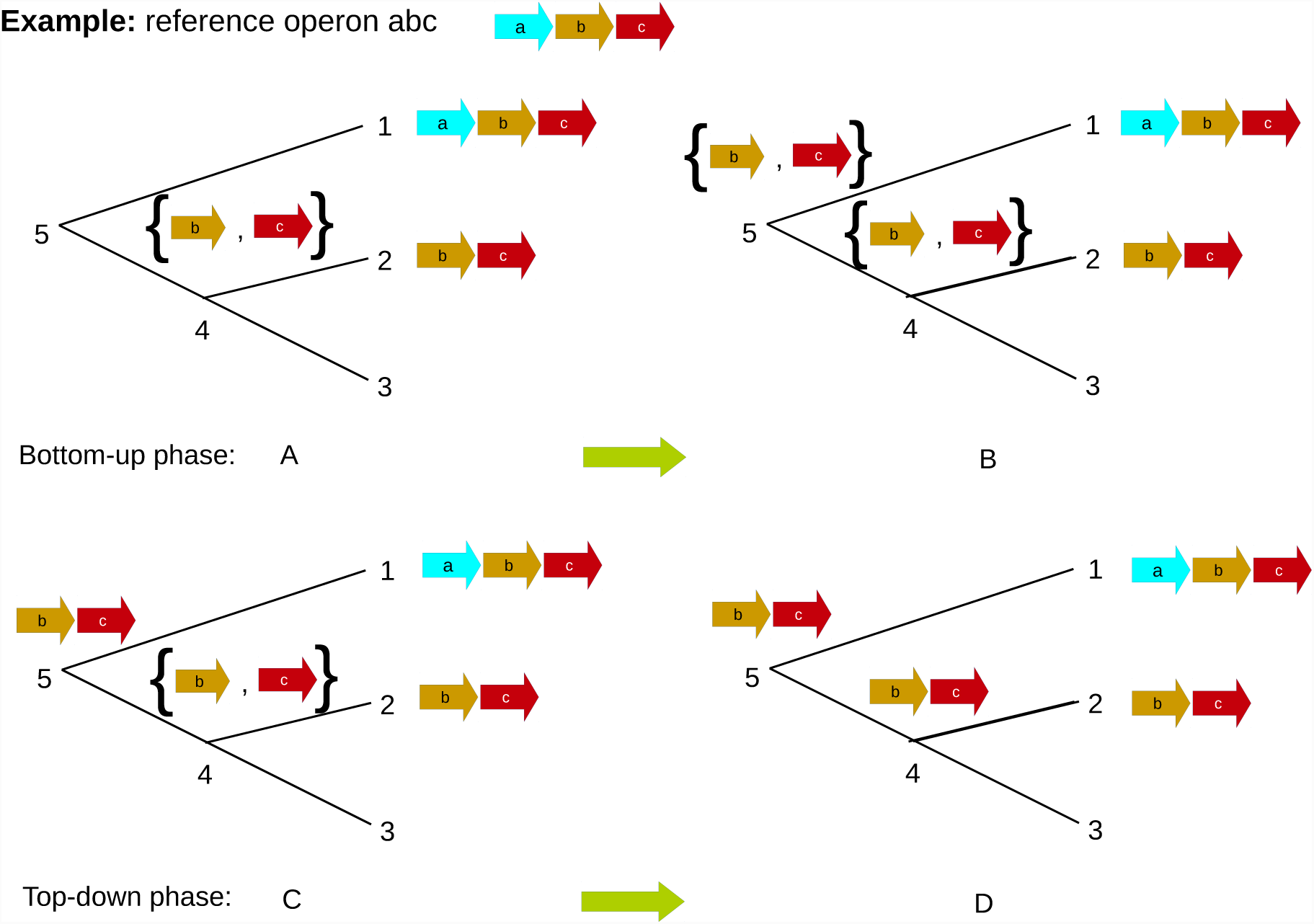
A simplified example of ancestral reconstruction using the global algorithm. In each panel, **1,2,3** are the extant nodes that are assigned with gene blocks **abc, bc,** ∅ respectively. The global algorithm traverses the tree bottom-up and top-down. In bottom up phase, the algorithm constructs the set of genes for the inner nodes (**4,5**). In panel A, at node **4**, the set of genes is {**b,c**} since **2,3** do not share any common genes. In panel B, at node **5**, the set of genes is {**b,c**} because **1,4** share genes **b,c**. In the top-down phase, the gene block is constructed for each inner node. In panel B, the gene block **bc** is assigned to node **5** using the set of genes of node **5** and the gene block of node **1**. We assign gene block **bc** to node **4** because of its set of genes and gene blocks in node **2**.

#### Algorithm 2: Global approach

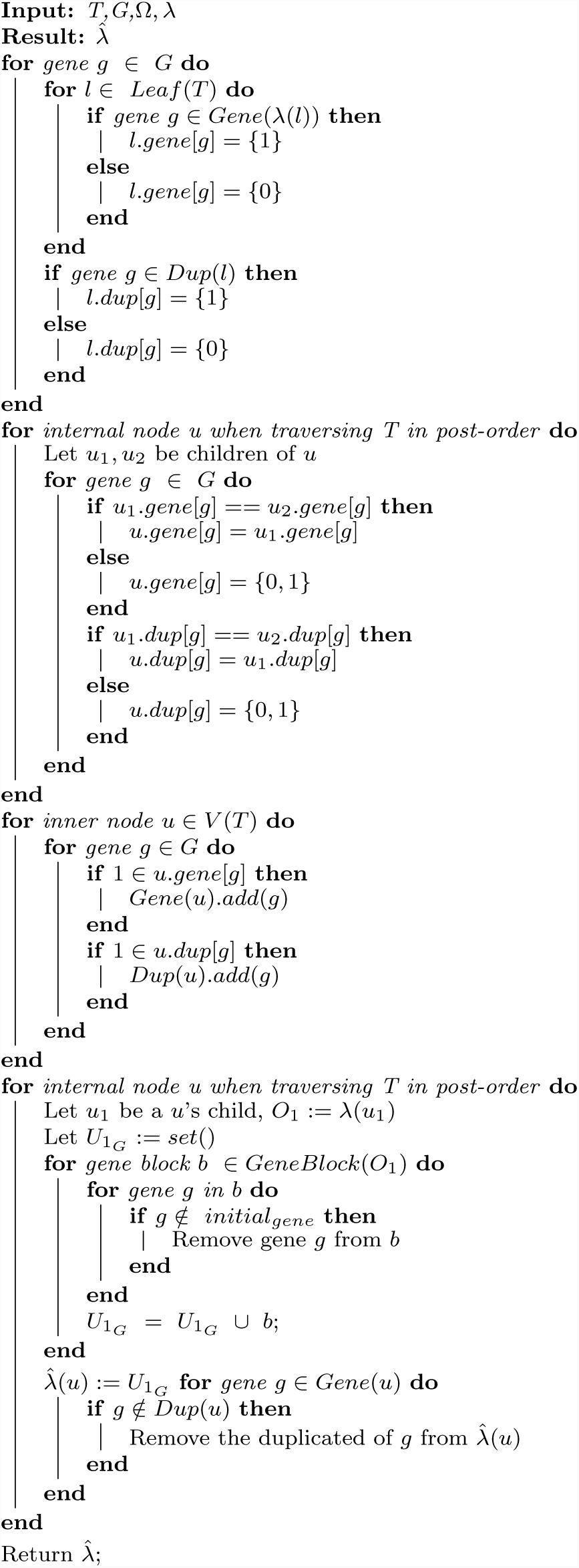

## Results and Discussion

We used experimentally-identified operons from *E. coli* K-12 and *B. subtilis* str. 168 genomes as gold standards for deriving operons from Gram-negative and Gram-positive bacteria, respectively. The reason we chose these two species is that they both have well-annotated genomes, including experimentally verified and functionally annotated operons.

### Operons from *Escherichia coli*

We chose *E. coli* as the reference species for proteobacteria, a major group of Gram-negative bacteria. Our selection resulted in a set of proteobacteria species comprising three *ϵ*-proteobacteria, six *α*-proteobacteria, seven *β*-proteobacteria and 17 *γ*-proteobacteria, including *E. coli*. These taxa include two *γ*-proteobacteria insect endosymbionts: *Buchnera aphidicola* and *Candidatus Blochmania*. These two species have unusually small genomes due to their endosymbiotic nature, and display massive gene loss. We reconstructed ancestors for the following operons from *E. coli* (described below): *atpIBEFHAGD* and *paaABCDEFGHIJK*.

#### atpIBEFHAGDC

The *atpIBEFHAGDC* operon codes for *F*_1_*F*_*o*_-ATPase, which catalyzes the synthesis of ATP from ADP and inorganic phosphate (Kasimoglu *et al.*, 1996). ATP synthase is composed of two fractions: F_1_ and F_o_ (Senior, 1990). The F_1_ fraction contains the catalytic sites and its proteins are coded by five genes (*atpA, atpC, atpD, atpG, atphH*) (Senior, 1990). The F_o_ complex constitutes the proton channel and its proteins are coded by three genes *atpF, atpE, atpB*. *atpI* is a non-essential regulatory gene. Figure S3 shows the high degree of conservation of this operon.

Figures 4 and 5 show ancestral reconstruction using the local and global maximum parsimony algorithms, respectively. Both local and global reconstructions show a consistency of having orthoblocks *atpACDGH* and *atpBF* in the most common ancestors for different Gram negative bacteria. This finding agrees with the long-standing hypothesis that the F_o_ and the F_1_ fractions have evolved separately, with the two fractions having homologs in the hexameric DNA helicases and with flagellar motor complexes. Although we find the gene *atpI* in several species, the reconstruction predicts that *atpI* is not in the same cluster with other genes. As stated, *atpI* is probably not an essential component of the *F*_1_*F*_*o*_ ATPase (NJ, 1984). Another interesting finding is the duplication of *atpF* in *E*-proteobacteria which appears to predate their common ancestor. Note that all genes exist as a gene block even in the endosymbionts *Blochmannia* and *B. aphidicola*.

**Fig 4.**
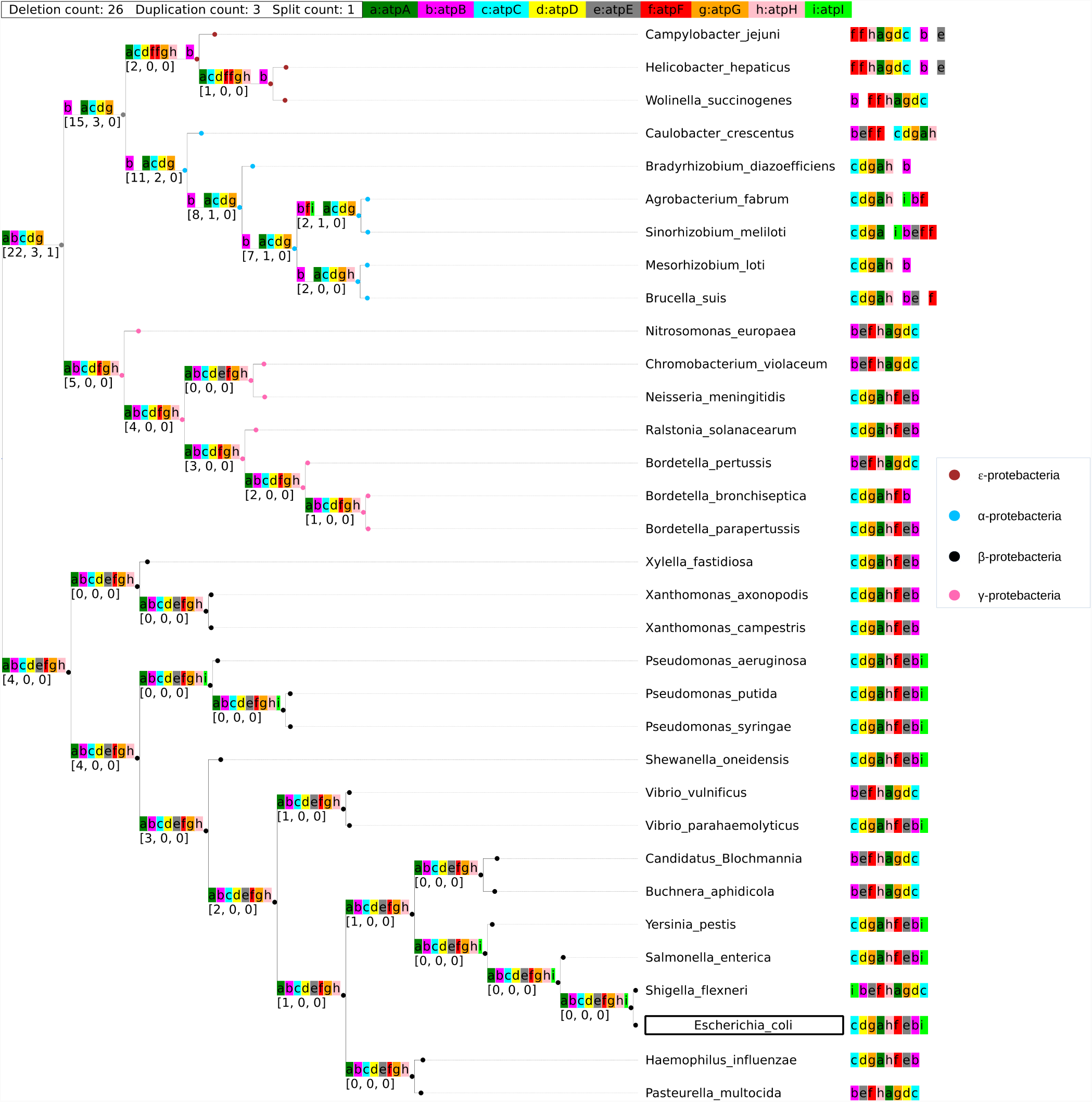
Ancestral reconstruction of operon *atpIBEFHAGDC* using the local optimization approach. The lower-case letters in each tree node represent the genes in the orthoblock (e.g. “a” represents “atpA”, see legend in blue bar, top). A ‘|’ designates a split (i.e. a distance *≥*500bp between the genes to either side of the ‘|’). The green bar on top shows the total number of events that took place in this reconstruction. The numbers in the brackets in the inner nodes are a 3-tuple showing the cumulative count of events going from the leafnodes to the tree root in the following order: [deletions, duplications, splits]. No orthologus gene blocks were found in species labeled with an asterisk (*). The reference genome E. coli is marked with a box. These naming and color conventions persist through this study.

**Fig 5.**
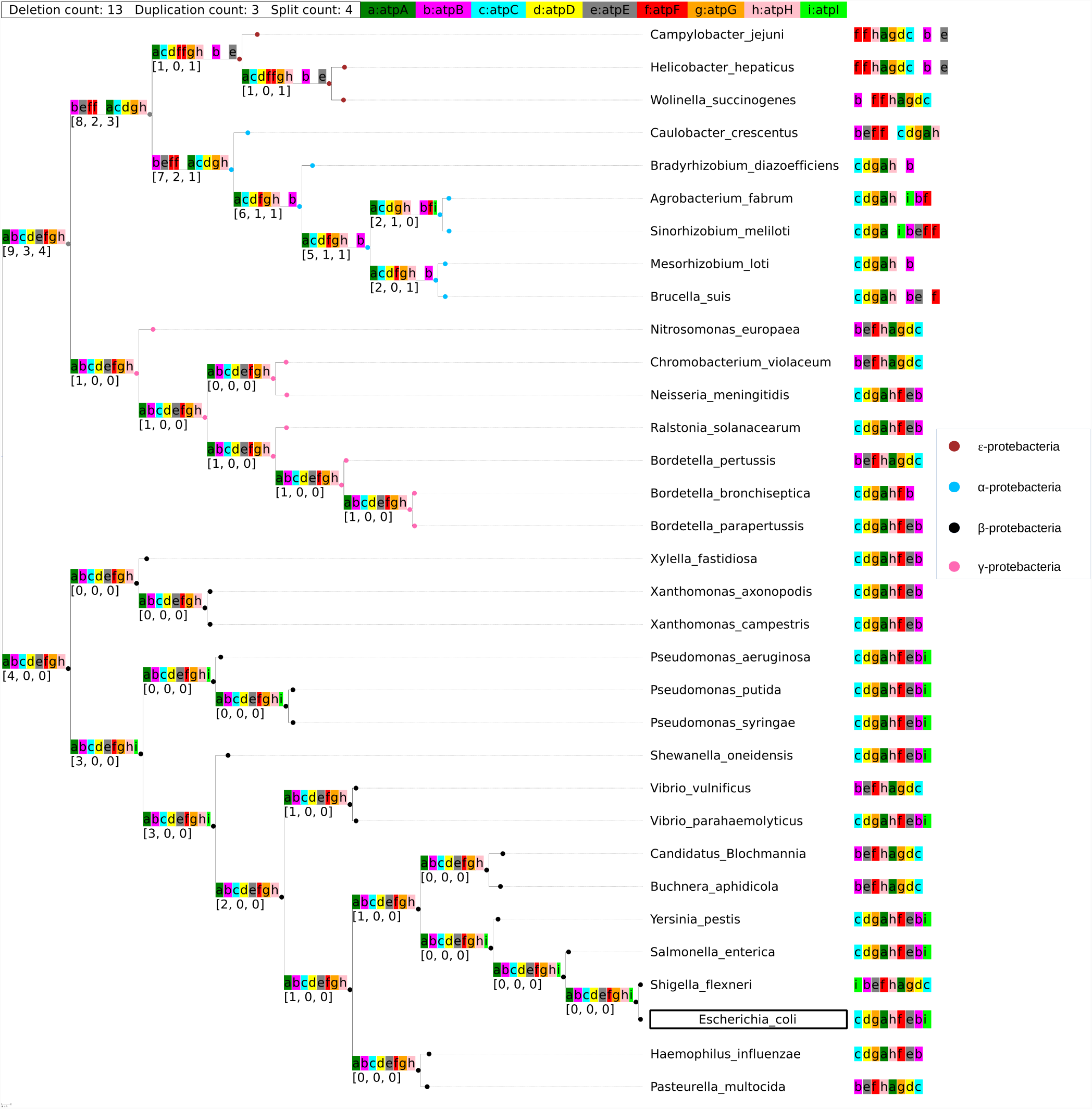
Ancestral reconstruction of operon *atpIBEFHAGDC* using the global optimization approach.

The *ϵ, α, β*, and *γ*-proteobacteria species all have a conserved intact F_1_ complex (coded by the *atpACDGH* cluster), which predates their common ancestor. The genes included in the F_o_ complex in *epsilon*-proteobacteria (gene products *atpB, atpE,atpF*) not in the same cluster as the genes making up F_1_. Furthermore, it is unclear whether the gene split that is only found in *ϵ*-proteobacteria is a split that predates the least common ancestor with the other proteobacteria clades, or whether it is a split introduced in the *E*-proteobacteria. From the reconstructions provided, the scenario appears to be the latter. Conversely, this observation may also be a result of the small number of species studied here. The species in the *ϵ* and *α*-proteobacteria display a known duplication of gene *atpF*. *atpF ′* appears as a sister group to *atpF* (Koumandou and Kossida, 2014).

#### paaABCDEFGHIJK

The operon *paaABCDEFGHIJK* codes for genes involved in the catabolism of phenylacetate (Martin and McInerney, 2009). The ability to catabolize phenylacetate varies greatly between proteobacterial species, and even among different *E. coli* K-12 strains. In contrast with *atpABCDEFG* operon which is conserved through many species, the operon *paaABCDEFGHIJK* is only found in full complement as an operon in some *E. coli* K-12 strains and some *Pseudomonas putida* strains. While obviously less conserved than the *atpABCDEFG* operon, certain orthoblocks appear to be conserved, providing possible partial functionality. The orthoblock *paaABCDE* is found in three *Bordetella* species and also in *Bradyrhizobium diazoefficiens*. The products of *paaA, paaB, paaC* and *paaE* make up the subunits of the 1,2-phenylacetyl-CoA epoxidase, and *paaD* is hypothesized to form an iron-sulfur cluster with the product of *paaE* (Grishin *et al.*, 2011). We did not find orthologs in the endosymbionts *B. aphidicola* and *Blochmannia*.

In both the local and global reconstructions (Figure S5 and Figure 6 respectively), only the ancestor of the *Bordetella* species have a combination of *paaABC* complex with *paaE*. It appears that only this combination has full activity (Grishin *et al.*, 2011). In addition, the global approach only predicts gene blocks for the ancestors of *α* and most of *γ*-proteobacteria. Only the common ancestor of the *Bordetella* genus contains the cluster *paaABCE*. It has been confirmed that this cluster of genes is identical to those of *E. coli* (Luengo, Garcia, and Olivera, 2001). In both approaches, gene *paaF* and *paaG* are not found to be in the same gene blocks, hence the ancestors are most likely missing the hydratase-isomerase complex. The *paaJ* thiolase catalyzes two steps in the phenylacetate catabolism (Ismail *et al.*, 2003; Teufel *et al.*, 2010; Nogales *et al.*, 2007). In addition, paaH is the NAD+-dependent 3-hydroxyadipyl-CoA dehydrogenase involved in phenylacetate catabolism(Ismail *et al.*, 2003). Therefore, it makes sense that *paaJ* and *paaH* appear in most of the ancestral nodes that have gene blocks.

**Fig 6.**
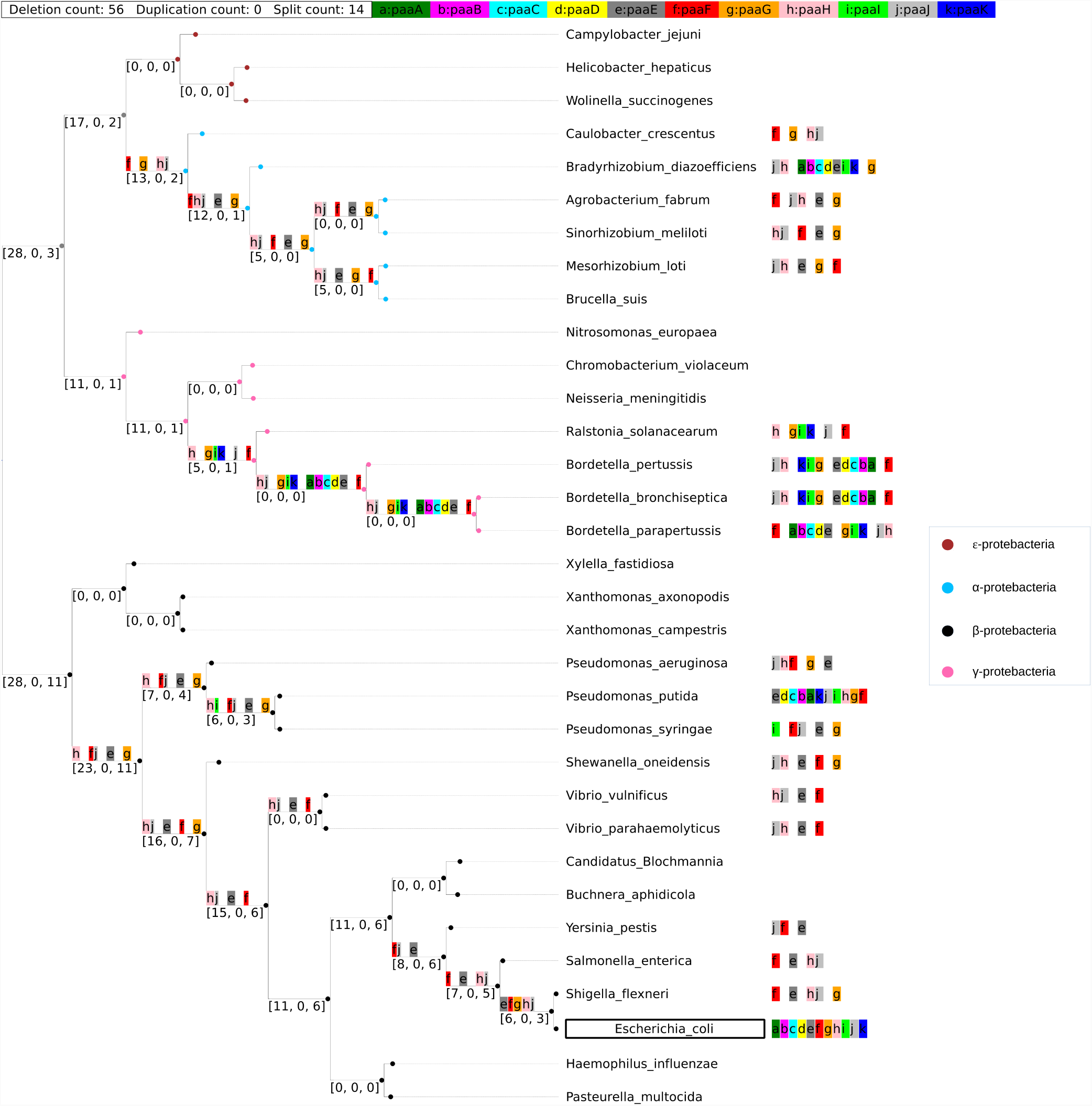
Ancestral gene block reconstruction of operon *paaABCDEFGHIJK* using the global reconstruction approach.

It is interesting to note that we see the formation of functional intermediate forms both in a highly conserved gene block *atpIBEFHAGDC* and the less conserved gene block based on the operon *paaABCDEFGHIJK*. Also, in both cases, the global approach performs better in term of minimizing events. For brevity, we only provide the global ancestral reconstruction henceforth.

### Operons from *Bacillus subtilis*

*B. subtilis* is a Gram-positive, spore forming bacterium commonly found in soil, and is also a normal gut commensal in humans. It is a model organism for Gram-positive spore forming bacteria, and as such its genome of about 4,450 genes is well annotated. Here we used ROAGUE to reconstruct the ancestors of two *B. subtilis* gene blocks across 33 species. We selected species from the order *Bacillales* using PDA. Species from the following families were selected: *Bacillaceae* (including the reference organism *B. subtilis*), *Staphylococcae*: macrococcus and staphylococcus, *Alicyclobacillaceae, Listeriaceae* and *Planococcaceae*.

#### lepA-hemN-hrcA-grpE-dnaK-dnaJ-prmA-yqeU-rimO

Gene block *lepA-hemN-hrcA-grpE-dnaK-dnaJ-prmA-yqeU-rimO* facilitates the heat shock response in *B. subtilis* and the gene block *hrcA-grpE-dnaK-dnaJ* was the first identified heat shock operon within *Bacillus spp* (Wetzstein *et al.*, 1992). The four genes *hrcA, grpE, dnaK, dnaJ* (e,c,b,a in Figure 7) form a tetracistronic structure, which is essential to the heat shock response role (Homuth *et al.*, 1997). The four genes are proximal (they never separated in the course of evolution) in all the species examined, and form the core of the orthoblock. Overall, this operon is quite conserved, and the ancestral reconstructions are highly similar to the reference operon.

**Fig 7.**
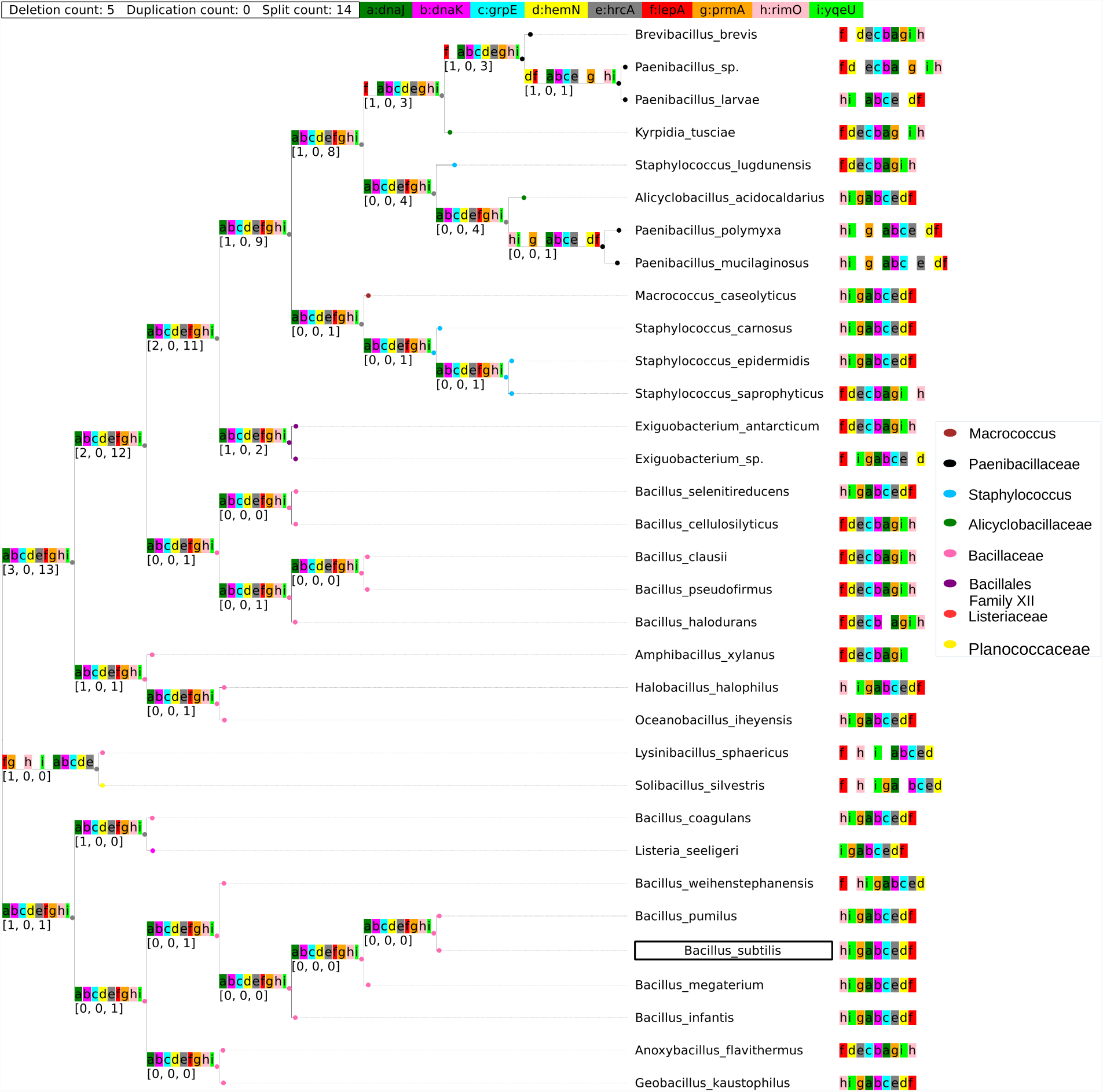
Ancestral reconstruction of *lepA-hemN-hrcA-grpE-dnaK-dnaJ-prmA-yqeU-rimO*.

#### mmgABCDE-prpB

The operon *mmgABCDE-prpB* is expressed during endosporulation (Acharya, 2009). Subunit *mmgABC*’s breakdown of fatty acids is a mean for attaining energy to drive the cell’s preparation for dormancy (Quattlebaum, 2009).Hence, it is reasonable to see that the common ancestor has this subunit. In addition, gene *mmgD* and gene *prpB/yqiQ* are predicted to be proximal. Several studies predicted that gene *mmgD, prpB*, and *prpD* encode the proteins of the putative methylcitrate shunt (Voigt *et al.*, 2007). However, they did not specify if deletion mutations might contribute to a defect of the functionality. See Figure 8.

**Fig 8.**
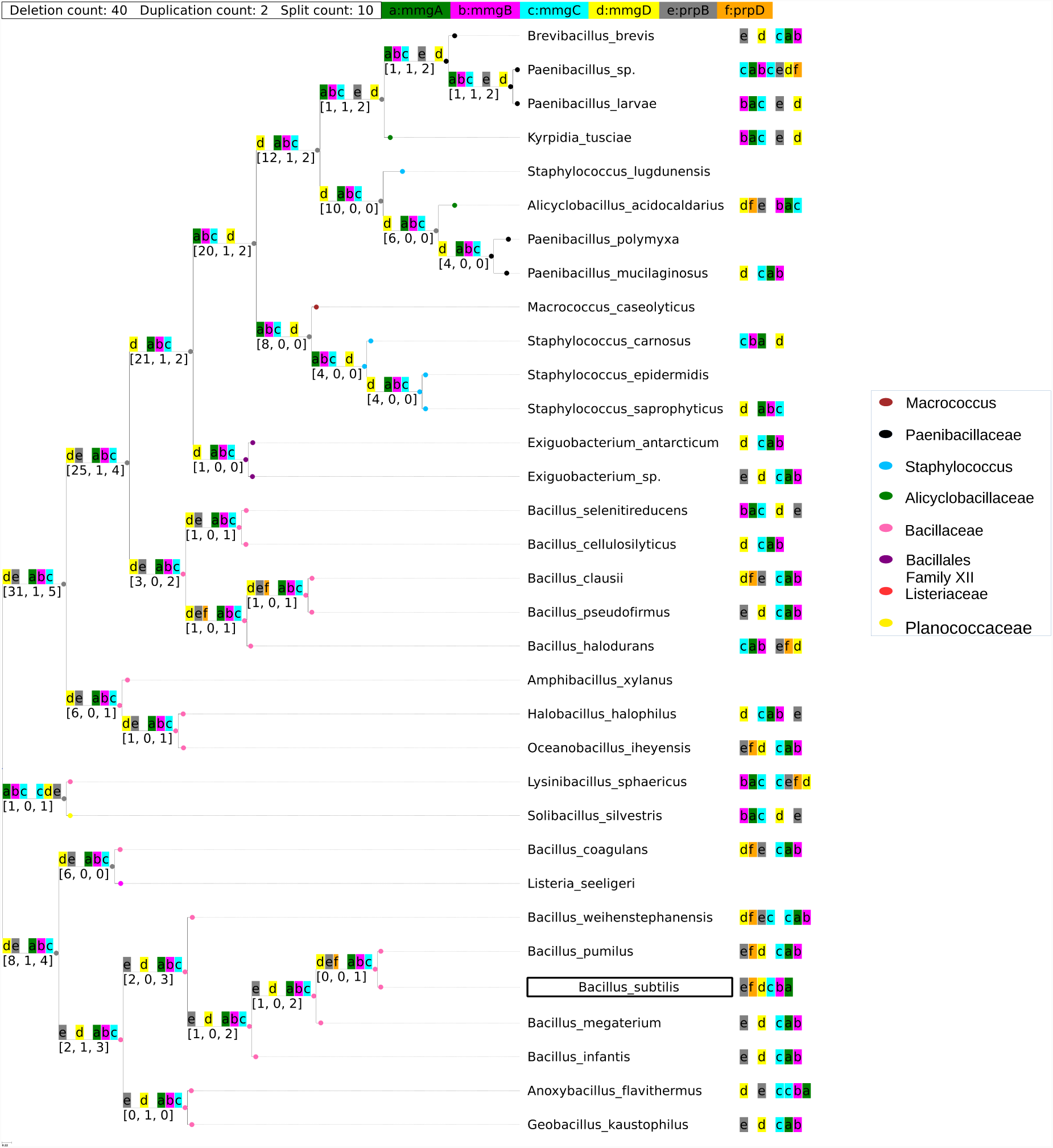
Ancestral reconstruction of *mmgABCDE-prpB*.

## Conclusions

Operons offer a tractable model for the evolution of complexity. Understanding how simple units of genes may converge into an operon can lead us to a better understanding of how a complex molecular systems evolve. Here we developed a method for to reconstruct ancestral gene blocks using maximum parsimony. Using this method we provide several examples of ancestral gene block reconstructions based on reference operons in *E. coli* and *B. subtilis*. Some interesting observations emerge regarding conservation and ancestry of operons. From our examples it appears that essentiality (the trait of being essential to life) and the formation of a protein complex are two drivers for gene block conservation. This is most apparent inthe *atpABCDEFG* operon coding for *F*_1_*F* _*o*_-atpase in proteobacteria. There are few evolutionary events identified in the *atpABCDEFG* operon ancestry. The ribose transporter block also seems to preserve the core ribose transporter (*rbsABC*), while not the ribose phosphorylation genes *rbsD* and *rbsK*. ROAGUE also highlights intermediate functional forms of the orthoblocks, as we see in the pattern of conservation in *paaABCDEFGHIJK*.

Our study does not account for horizontal gene transfer, which has been shown to be major driver in operon dispersal in distant species (Omelchenko *et al.*, 2003; Koonin, 2009). Detecting horizontal gene transfer is typically done by looking for conservation of genes and gene structures between distant OTUs, and for anomalous codon usage (Koonin, Makarova, and Aravind, 2001). Our method opens up a new way of HGT detection, by reconciling a species tree with an operon tree, in the same waythat phylogenomic analyses dofor gene trees and species trees (Eisen, 1998), which would be an interesting future development of this study. In addition, we ignore the gene order in the gene block. While the relationship between gene organization and expression in operons is not well understood, it is clear from several studies that gene order does have an effect on expression and on the functionality of the operon in general (e.g. (Hiroe *et al.*, 2012; Wells, Bergendahl, and Marsh, 2016; Lim, Lee, and Hussein, 2011)). Adding the parameters of horizontal gene transfer, gene order preservation, or both to ROAGUE would be highly valuable. Weinvite the communityto contribute to ROAGUE, as well as use the tool for identifying orthologous gene blocks, and reconstructing their ancestry.

## References

Acharya, R (2009). “Overexpression, purification, and characterization of MmgD from Bacillus subtilis strain 168”. PhD thesis. University of North Carolina at Greensboro.

Adami, C, Ofria, C, and Collier, TC (2000). “Evolution of biological complexity”. Proceedings of the National Academy of Sciences 97.9, pp. 4463–4468.

Alm, E, Huang, K, and Arkin, A (2006). “The Evolution of Two-Component Systems in Bacteria Reveals Different Strategies for Niche Adaptation”. PLOS Computational Biology 2.11, e143+.

Bonner, JT (1988). The Evolution of Complexity by Means of Natural Selection. Princeton University Press.

Bush, EC et al. (2018). “xenoGI: reconstructing the history of genomic island insertions in clades of closely related bacteria.” BMC bioinformatics 19.1.

Chernomor, O et al. (2015). “Split diversity in constrained conservation prioritization using integer linear programming”. Methods in Ecology and Evolution 6.1, pp. 83–91.

Eisen, JA (1998). “A phylogenomic study of the MutS family of proteins.” Nucleic acids research 26.18, pp. 4291–4300.

Faith, DP (1992). “Conservation evaluation and phylogenetic diversity”. Biological conservation 61.1, pp. 1–10.

Fani, R, Brilli, M, and Lio, P (2005). “The origin and evolution of operons: the piecewise building of the proteobacterial histidine operon.” Journal of molecular evolution 60.3, pp. 378–390.

Goldberg, T, Rost, B, and Bromberg, Y (2016). “Computational prediction shines light on type III secretion origins.” Scientific reports 6.

Grishin, AM et al. (2011). “Structural and Functional Studies of the Escherichia coli Phenylacetyl-CoA Monooxygenase Complex”. Journal of Biological Chemistry 286.12, pp. 10735–10743.

Hiroe, A et al. (2012). “Rearrangement of Gene Order in the phaCAB Operon Leads to Effective Production of Ultrahigh-Molecular-Weight Poly[(R)-3-Hydroxybutyrate] in Genetically Engineered Escherichia coli”. Applied and Environmental Microbiology 78.9, pp. 3177–3184.

Homuth, G et al. (1997). “The dnaK operon of Bacillus subtilis is heptacistronic.” Journal of bacteriology 179.4, pp. 1153–1164.

Horowitz, NH (1945). “On the Evolution of Biochemical Syntheses.” Proceedings of the National Academy of Sciences of the United States of America 31.6, pp. 153–157.

Hsiao, WW et al. (2005). “Evidence of a large novel gene pool associated with prokaryotic genomic islands.” PLoS genetics 1.5, e62+.

Ismail, W et al. (2003). “Functional genomics by NMR spectroscopy”. European Journal of Biochemistry 270.14, pp. 3047–3054.

Kasimoglu, E et al. (1996). “Transcriptional regulation of the proton-translocating ATPase (atpIBEFHAGDC) operon of Escherichia coli: control by cell growth rate.” Journal of bacteriology 178.19, pp. 5563–5567.

Koonin, EV, Makarova, KS, and Aravind, L (2001). “Horizontal gene transfer in prokaryotes: quantification and classification.” Annual review of microbiology 55.1, pp. 709–742.

Koonin, EV (2009). “Evolution of genome architecture.” The international journal of biochemistry & cell biology 41.2, pp. 298–306.

Koonin, EV and Dolja, VV (2006). “Evolution of complexity in the viral world: the dawn of a new vision.” Virus research 117.1, pp. 1–4.

Koumandou, VL and Kossida, S (2014). “Evolution of the F 0 F 1 ATP synthase complex in light of the patchy distribution of different bioenergetic pathways across prokaryotes”. PLoS Comput Biol 10.9, e1003821.

Lawrence, JG and Roth, JR (1996). “Selfish operons: horizontal transfer may drive the evolution of gene clusters.” Genetics 143.4, pp. 1843–1860.

Lim, HN, Lee, Y, and Hussein, R (2011). “Fundamental relationship between operon organization and gene expression”. Proceedings of the National Academy of Sciences 108.26, pp. 10626–10631.

Luengo, JM, Garcia, JL, and Olivera, ER (2001). “The phenylacetyl-CoA catabolon: a complex catabolic unit with broad biotechnological applications”. Molecular microbiology 39.6, pp. 1434–1442.

Lynch, M and Conery, JS (2003). “The Origins of Genome Complexity”. Science 302.5649, pp. 1401–1404.

Martin, FJ and McInerney, JO (2009). “Recurring cluster and operon assembly for phenylacetate degradation genes”. BMC evolutionary biology 9.1, p. 36.

Nj, G (1984). “Construction and characterization of an Escherichia coli strain with a uncI mutation.” Journal of Bacteriology 158, pp. 820–825.

Nogales, J et al. (2007). “Characterization of the last step of the aerobic phenylacetic acid degradation pathway”. Microbiology 153.2, pp. 357–365.

Omelchenko, M et al. (2003). “Evolution of mosaic operons by horizontal gene transfer and gene displacement in situ”. Genome Biology 4.9, R55+.

Pál, C and Papp, B (2017). “Evolution of complex adaptations in molecular systems.” Nature ecology & evolution 1.8, pp. 1084–1092.

Price, MN, Arkin, AP, and Alm, EJ (2006). “The Life-Cycle of Operons”. PLOS Genetics 2.6, e96+.

Quattlebaum, AL (2009). “Characterization of Biosynthetic and Catabolic Pathways of Bacillus Subtilis Strain 168”. PhD thesis. University of North Carolina at Greensboro.

Ream, DC, Bankapur, AR, and Friedberg, I (2015). “An event-driven approach for studying gene block evolution in bacteria”. Bioinformatics 31.13, pp. 2075–2083.

Senior, AE (1990). “The Proton-Translocating ATPase of Escherichia Coli”. Annual Review of Biophysics and Biophysical Chemistry 19.1. PMID: 2141983, pp. 7–41.

Stahl, FW and Murray, NE (1966). “The evolution of gene clusters and genetic circularity in microorganisms.” Genetics 53.3, pp. 569–576.

Teufel, R et al. (2010). “Bacterial phenylalanine and phenylacetate catabolic pathway revealed”. Proceedings of the National Academy of Sciences 107.32, pp. 14390–14395.

Voigt, B et al. (2007). “The glucose and nitrogen starvation response of Bacillus licheniformis”. Proteomics 7.3, pp. 413–423.

Wagner, GP and Altenberg, L (1996). “Perspective: Complex Adaptations and the Evolution of Evolvability”. Evolution 50.3, pp. 967–976.

Wells, JN, Bergendahl, LT, and Marsh, JA (2016). “Operon Gene Order Is Optimized for Ordered Protein Complex Assembly.” Cell reports 14.4, pp. 679–685.

Wetzstein, M et al. (1992). “Cloning, sequencing, and molecular analysis of the dnaK locus from Bacillus subtilis.” Journal of Bacteriology 174.10, pp. 3300–3310.

