## Supplemental Material for "Tracing the Ancestry of Operons in Bacteria"

### Supplementary

This section contains analysis of two additional gene blocks, the correctness and proof of the two algorithms discussed in this work. In the end, we also include the two phylomatrix of gene blocks *atpIBEFHAGDC* and *paaABCDEFGHIJK*, and the local reconstruction of gene block *paaABCDEFGHIJK*.

#### Additional Gene Blocks

Here, we provide another ancestral reconstructions of two more gene blocks in *E. coli*.

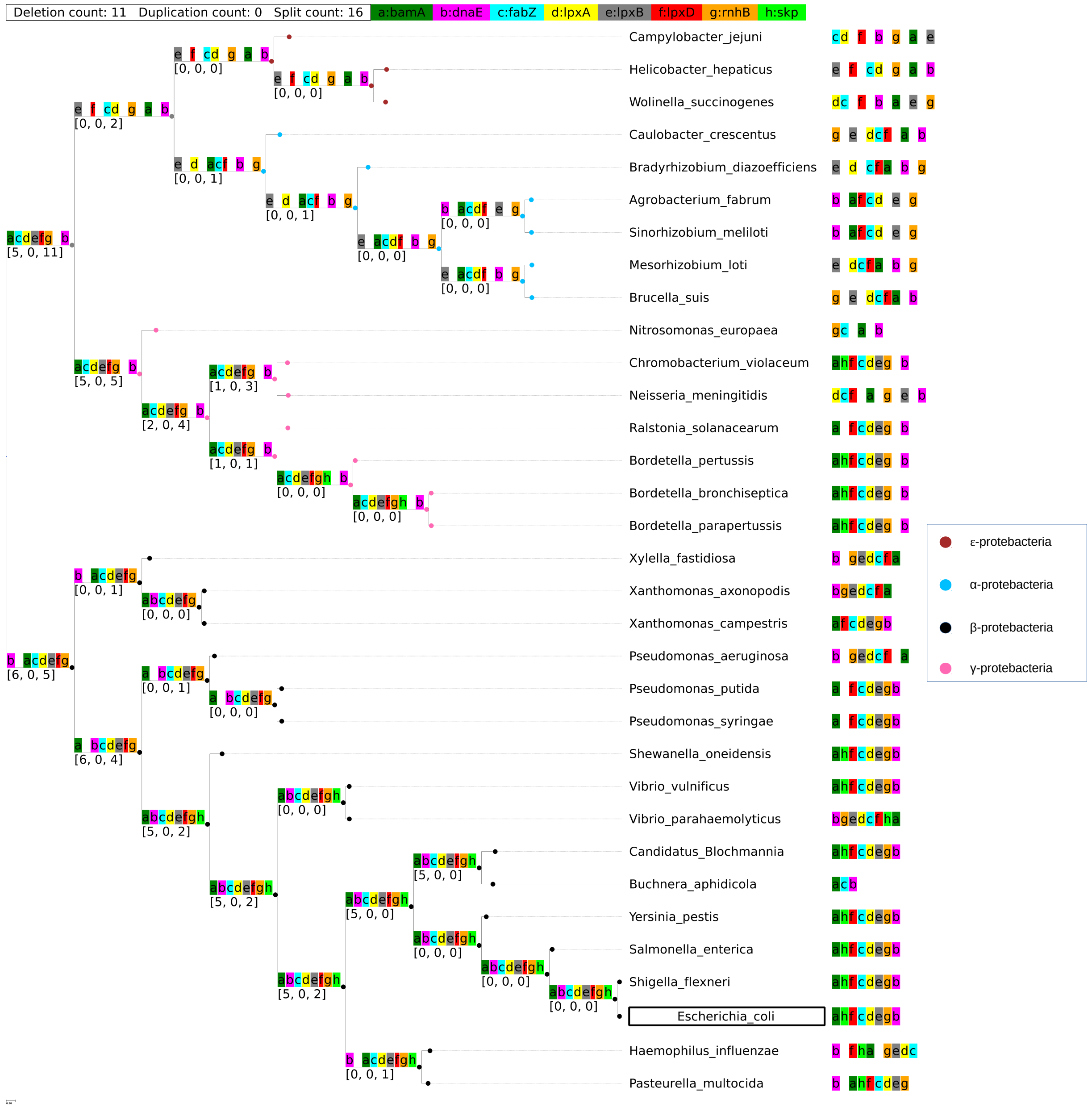


**Fig S1.** Ancestral reconstrucion of gene block *bamA-skp-lpxD-fabZ-lpxAB-rnhB-dnaE*

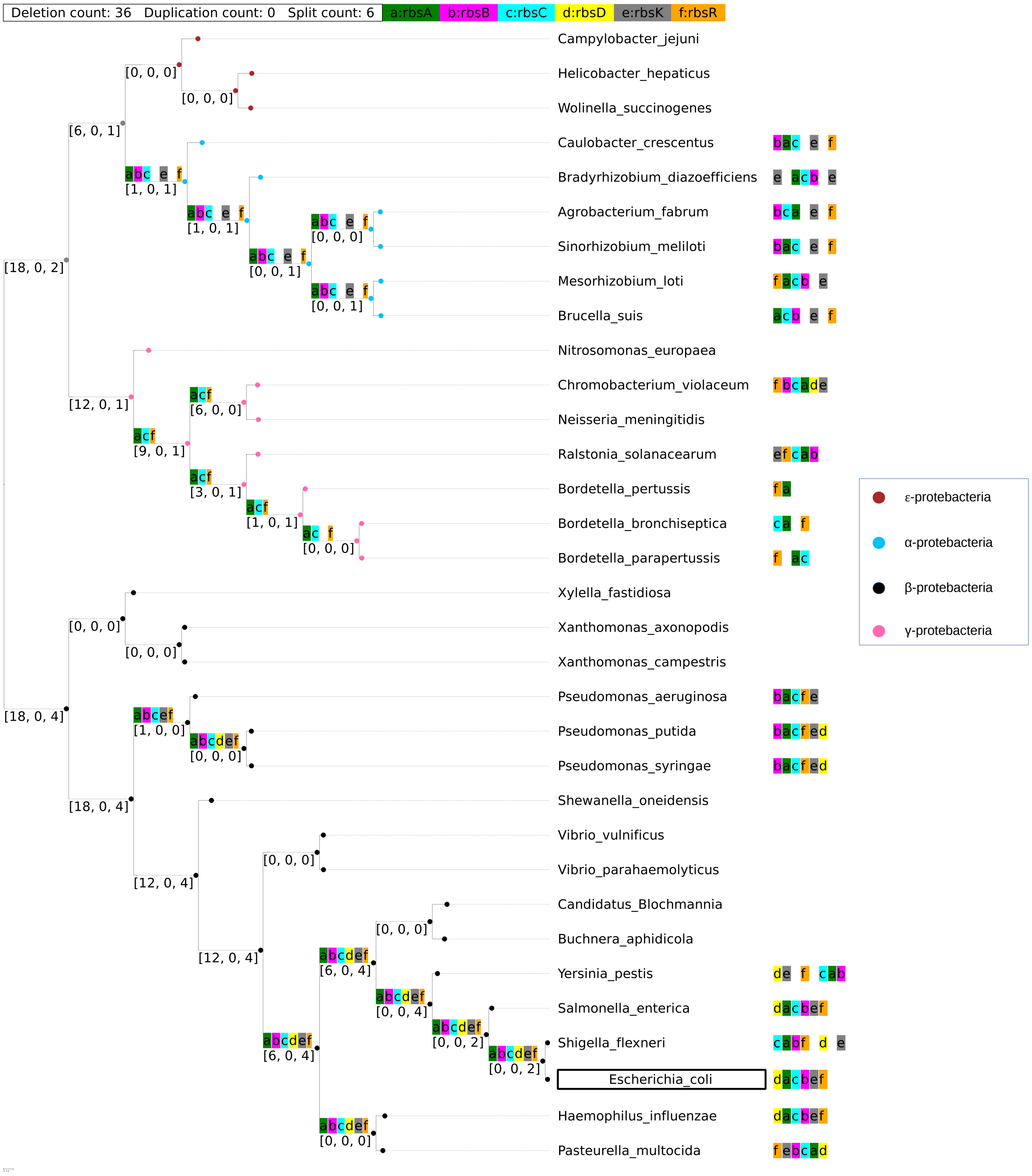


**Fig S2.** Ancestral reconstruction of *rbsDACBKR*. .

***bamA-skp-lpxD-fabZ-lpxAB-rnhB-dnaE***. The operon *bamA-skp-lpxD-fabZ-lpxAB-rnhB-dnaE* participates in DNA replication, repair, immune reaction, and signal transduction. It is actually a complex regulon with several promoter sites Liang and Liu, 2008. Gene *bamA* is highly conserved Gentle *et al.*, 2004 and is required for Gram-negative outer membrane protein assembly Doerrler and Raetz, 2005; Werner and Misra, 2005. Gene *dnaE* encodes the alpha-catalytic subunit of the DNA polymerase III holoenzyme Maki and Kornberg, 1985. The reconstruction result has shown that those two genes have appeared in all the ancestors. Note that *bamA* is predicted to not be in the same regulatory block as the rest of the operon in

$\gamma$-proteobacteria. At the same time, gene *dnaE* is not in the same block of the operon in $\beta$-proteobacteria. However, these two splits should not affect the overall operon functionality since neither *bamA* nor *dnaE* are found to form a subunit with another gene in the operon. At the same time, the cluster of *lpxD-fabZ-lpxA* is involved in lipid A biosynthesis in many bacteriaSchmid *et al.*, 1989; Mohan *et al.*, 1994.

***rbsDACBKR***. The operon *rbsDACBKR* expresses genes associated with the ribose transport complex in *E. coli* Zaitseva *et al.*, 1996; Barroga *et al.*, 1996. The *rbsABC* genes compose an ATP-dependent ribose transporter that is a member of the ATP-Binding Cassette (ABC) superfamily of transporters Park and Park, 1999. Mutations in each of the components eliminated transport of ribose at an external concentration of 1$\mu$M, indicating that the components make up a transport system that is responsible for high-affinity ribose transport Iida *et al.*, 1984. From the reconstruction, we observe that the core gene cluster of the transporter *rbsABC* starts forming in three different inner nodes: (1) the common ancestor of $\alpha$-proteobacteria; (2) $\gamma$-proteobacteria (genus *Pseudomonas*), and (3) $\gamma$-proteobacteria (*Enterobacteriaceae, Pasteurellaceae* families). The three other genes, *rbsK*, *rbsD* and *rbsR* are not essential for ribose transport. *rbsR* codes for the repressor protein which regulates the operon Shimada, Kori, and Ishihama, 2013; Mauzy and Hermodson, 1992. *rbsD*, and *rbsK* are involved in the conversion of D-ribose to D-ribose 5-phosphate Oh, Park, and Park, 1999. The gene block is most complete in the $\gamma$-proteobacteria, but the core transport genes appear also at the common ancestors of the $\alpha$-proteobacteria.

#### Correctness and Proof: Local Optimum

##### Correctness

Let $\hat{\lambda} := Algorithm\ 1(T, G, \Omega, \lambda)$. For each $u \in I(T)$, let $u_1, u_2$ be its children. Let $O, O_1, O_2$ respectively be the orthoblock assigned to $u, u_1, u_2$ by function $\hat{\lambda}$. We will show that our results minimize $d_d(O, O_1) + d_d(O, O_2)$ and $d_u(O, O_1) + d_u(O, O_2)$

**Lemma 1**: $\forall g \in G$, if $FREQ_g(u) > .5$ then either $FREQ_g(u_1) > .5$ or $FREQ_g(u_2) > .5$ In addition, if $FREQ_g(u) \leq .5$ then either $FREQ_g(u_1) \leq .5$ or $FREQ_g(u_2) \leq .5$
*Proof*:

1. If $FREQ_g(u) > .5$ then either $FREQ_g(u_1) > .5$ or $FREQ_g(u_2) > .5$
   Assume that $FREQ_g(u_1) \leq .5$ and $FREQ_g(u_2) \leq .5$, then

   $$\begin{cases} |\{v \in HasLeaf(u_1) | g \in Gene(\lambda(v)\}| \leq \frac{|HasLeaf(u_1)|}{2} \\ |\{v \in HasLeaf(u_2) | g \in Gene(\lambda(v)\}| \leq \frac{|HasLeaf(u_2)|}{2} \end{cases}$$

   Define $H := \{v \in (HasLeaf(u_1) \cup HasLeaf(u_2)) | g \in Gene(\lambda(v)\}$, from the two inequalities above, we have:

   $$\left|H\right| \leq \frac{|HasLeaf(u_1)|}{2} + \frac{|HasLeaf(u_2)|}{2}$$

   Since $u_1, u_2$ are the children of $u$, then

   $$\begin{cases} HasLeaf(u_1) \cup HasLeaf(u_2) = HasLeaf(u) \\ HasLeaf(u_1) \cap HasLeaf(u_2) = \emptyset \end{cases}$$

   $$\rightarrow \left|\{v \in HasLeaf(u) | g \in Gene(\lambda(v)\}\right| \leq \frac{|HasLeaf(u)|}{2}$$
   $$\rightarrow FREQ_g(u) \leq .5$$

   By contraposition, if $FREQ_g(u) > .5$ then either $FREQ_g(u_1) > .5$ or $FREQ_g(u_2) > .5$

2. If $FREQ_g(u) \leq .5$ then either $FREQ_g(u_1) \leq .5$ or $FREQ_g(u_2) \leq .5$
   We can prove it using the same logic as above.

**Lemma 2**: $\forall g \in G$, if $g \in Gene(O)$ *and* $g \notin Gene(O')$, then $|I_g(O) - I_g(O_1)| + |I_g(O) - I_g(O_2)| \leq |I_g(O') - I_g(O_1)| + |I_g(O') - I_g(O_2)|$ .
*Proof*:
Since $g \in Gene(O)$, then $FREQ_g(u) > .5$. Therefore, $FREQ_g(u_1) > .5$ *or* $FREQ_g(u_2) > .5$ (by lemma 1). Hence, $g \in Gene(u_1)$ *or* $g \in Gene(u_2)$. Consider 3 cases:

1. If $u_1$ and $u_2$ contain $g$, then
   $|I_g(O) - I_g(O_1)| + |I_g(O) - I_g(O_2)| = |1 - 1| + |1 - 1| = 0$
   $|I_g(O') - I_g(O_1)| + |I_g(O') - I_g(O_2)| = |0 - 1| + |0 - 1| = 2$
   Therefore, $|I_g(O) - I_g(O_1)| + |I_g(O) - I_g(O_2)| < |I_g(O') - I_g(O_1)| + |I_g(O') - I_g(O_2)|$

2. If only $u_1$ contains $g$, then
   $|I_g(O) - I_g(O_1)| + |I_g(O) - I_g(O_2)| = |1 - 1| + |1 - 0| = 1$
   $|I_g(O') - I_g(O_1)| + |I_g(O') - I_g(O_2)| = |0 - 1| + |0 - 0| = 1$
   Therefore, $|I_g(O) - I_g(O_1)| + |I_g(O) - I_g(O_2)| = |I_g(O') - I_g(O_1)| + |I_g(O') - I_g(O_2)|$

3. If only $u_2$ contains $g$, then
   $|I_g(O) - I_g(O_1)| + |I_g(O) - I_g(O_2)| = |1 - 0| + |1 - 1| = 1$
   $|I_g(O') - I_g(O_1)| + |I_g(O') - I_g(O_2)| = |0 - 0| + |0 - 1| = 1$
   Therefore, $|I_g(O) - I_g(O_1)| + |I_g(O) - I_g(O_2)| = |I_g(O') - I_g(O_1)| + |I_g(O') - I_g(O_2)|$

From the above cases, we conclude that $|I_g(O)-I_g(O_1)|+|I_g(O)-I_g(O_2)| \leq |I_g(O')-I_g(O_1)|+|I_g(O')-I_g(O_2)|$

**Lemma 3**: $\forall g \in G$, if $g \notin Gene(O)$ *and* $g \in Gene(O')$, then $|I_g(O) - I_g(O_1)| + |I_g(O) - I_g(O_2)| \leq |I_g(O') - I_g(O_1)| + |I_g(O') - I_g(O_2)|$ .
*Proof*:
Since $g \notin Gene(O)$, then $FREQ_g(u) < .5$. Therefore, $FREQ_g(u_1) < .5$ *or* $FREQ_g(u_2) < .5$ (by lemma 1. Hence, $g \notin Gene(u_1)$ *or* $g \notin Gene(u_2)$. Consider 3 cases:

1. If $u_1$ and $u_2$ do not contain $g$, then
$|I_g(O) - I_g(O_1)| + |I_g(O) - I_g(O_2)| = |0 - 0| + |0 - 0| = 0$
$|I_g(O') - I_g(O_1)| + |I_g(O') - I_g(O_2)| = |1 - 0| + |1 - 0| = 2$
Therefore, $|I_g(O) - I_g(O_1)| + |I_g(O) - I_g(O_2)| < |I_g(O') - I_g(O_1)| + |I_g(O') - I_g(O_2)|$

2. If only $u_1$ does not contain $g$, then
$|I_g(O) - I_g(O_1)| + |I_g(O) - I_g(O_2)| = |0 - 0| + |0 - 1| = 1$
$|I_g(O') - I_g(O_1)| + |I_g(O') - I_g(O_2)| = |1 - 0| + |1 - 1| = 1$
Therefore, $|I_g(O) - I_g(O_1)| + |I_g(O) - I_g(O_2)| = |I_g(O') - I_g(O_1)| + |I_g(O') - I_g(O_2)|$

3. If only $u_2$ does not contain $g$, then
$|I_g(O) - I_g(O_1)| + |I_g(O) - I_g(O_2)| = |1 - 1| + |1 - 0| = 1$
$|I_g(O') - I_g(O_1)| + |I_g(O') - I_g(O_2)| = |0 - 1| + |0 - 0| = 1$
Therefore, $|I_g(O) - I_g(O_1)| + |I_g(O) - I_g(O_2)| = |I_g(O') - I_g(O_1)| + |I_g(O') - I_g(O_2)|$

From the above cases, we conclude that $|I_g(O)-I_g(O_1)|+|I_g(O)-I_g(O_2)| \leq |I_g(O')-I_g(O_1)|+|I_g(O')-I_g(O_2)|$

1. Minimal deletions: Given an assignment of orthoblock $O'$ to $u$, we will show that $d_d(O', O_1) + d_d(O', O_2) \geq d_d(O, O_1) + d_d(O, O_2)$

   *Proof*:

$$
\begin{aligned}
d_d(O', O_1) + d_d(O', O_2) &= \sum_g (|I_g(O') - I_g(O_1)| + |I_g(O') - I_g(O_2)|) \\
&= \sum_{g \in O'} (|I_g(O') - I_g(O_1)| + |I_g(O') - I_g(O_2)|) + \\
&\sum_{g \notin O'} (|I_g(O') - I_g(O_1)| + |I_g(O') - I_g(O_2)|) \\
&= \sum_{g \in O', g \in O} (|I_g(O') - I_g(O_1)| + |I_g(O') - I_g(O_2)|) + \\
&\sum_{g \in O', g \notin O} (|I_g(O') - I_g(O_1)| + |I_g(O') - I_g(O_2)|) + \\
&\sum_{g \notin O', g \in O} (|I_g(O') - I_g(O_1)| + |I_g(O') - I_g(O_2)|) + \\
&\sum_{g \notin O', g \notin O} (|I_g(O') - I_g(O_1)| + |I_g(O') - I_g(O_2)|) \\
&\geq \sum_{g \in O', g \in O} (|I_g(O) - I_g(O_1)| + |I_g(O) - I_g(O_2)|) + \\
&\sum_{g \in O', g \notin O} (|I_g(O) - I_g(O_1)| + |I_g(O) - I_g(O_2)|) + \\
&\sum_{g \notin O', g \in O} (|I_g(O) - I_g(O_1)| + |I_g(O) - I_g(O_2)|) + \\
&\sum_{g \notin O', g \notin O} (|I_g(O) - I_g(O_1)| + |I_g(O) - I_g(O_2)|) \\
&= \sum_{g \in O} (|I_g(O) - I_g(O_1)| + |I_g(O) - I_g(O_2)|) + \\
&\sum_{g \notin O} (|I_g(O) - I_g(O_1)| + |I_g(O) - I_g(O_2)|) \\
&= d_d(O, O_1) + d_d(O, O_2)
\end{aligned} \tag{1}
$$

2. Minimal duplication:

   *Proof:*

   Applying the same idea as the above proof with $DUP_g(u), Dup(u)$ instead of $FREQ_g(u), Gene(u)$ , we will achieve same result.

##### Runtime

The main challenge is how to store the the data of $FREQ_g(v), HasLeaf(v)$ for each inner node $v$. This can be done with dynamic programming. Algorithm 1 runs in polynomial time. Together, the algorithm takes $O(m^2) \times O(n) = O(m^2 \times n)$ with $n$ is the number of leaf nodes, and $m$ as the number of genes in the reference orthoblock.

#### Correctness and Proof: Global Optimum

##### Correctness

Let $\hat{\lambda} := Algorithm\ 2(T, G, \Omega, \lambda)$. We will show that $d_d(\hat{\lambda}) := \sum_{(u,v) \in E}(d_d(u, v))$ and $d_u(\hat{\lambda}) := \sum_{(u,v) \in E}(d_u(u, v))$ are minimal.

1. Minimal deletions:

   As stated above, $d_d(O,O') := |\sum_g (I_g(O) - I_g(O'))|$. Therefore, we can rewrite out global deletion cost as:

$$d_d(\hat{\lambda}) := \sum_{(u,v)\in E} (d_d(u,v)) = \sum_{(u,v)\in E} (|\sum_g (I_g(\hat{\lambda}(u)) - I_g(\hat{\lambda}(v)))|)$$

   Since each gene occurrence within a gene block is independent from each other, we only need to show that our algorithm provide a global minimum deletion for any genes $g$. Our algorithm is based on Fitch algorithm, and the proof can be followed by the conventional proof of Fitch easily.

2. Minimal duplications:

   *Proof:*

   Applying the same idea as the above proof with $DUP_g(u), Dup(u)$ instead of $FREQ_g(u), Gene(u)$ , we will achieve same result.

#### Run Time

This algorithm is twice as slow as the Local Algorithm. The reason is that it has to traverse the tree twice, in post order and level order. However,it still takes $O(m^2 \times n)$ to finish.

#### Phylomatrices

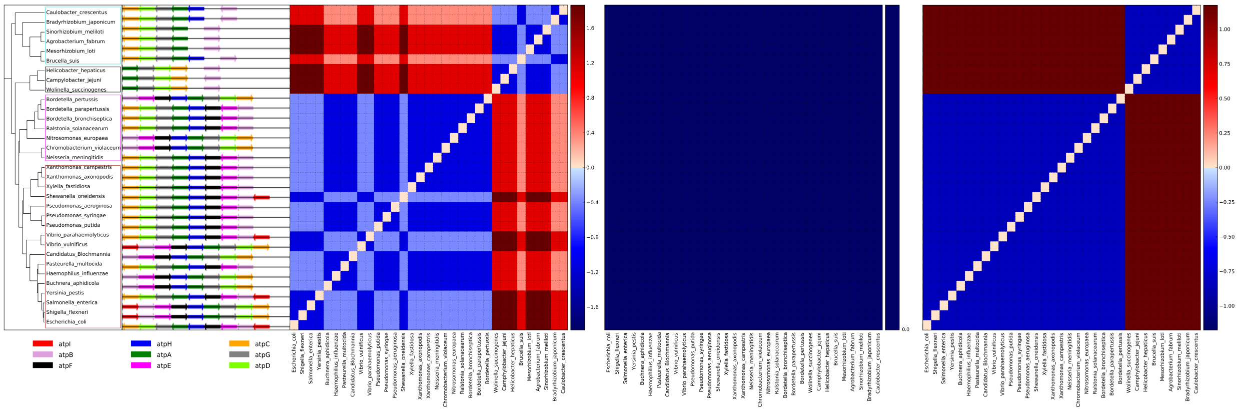


**Fig S3.** A phylomatrix of gene block *atpIBEFHAGDC*. Each matrix cell depicts the degree of relative conservation (less likely to happen) of the event between any two species. Blue is more conserved, red is less conserved. Squares, left to right: conservation of deletions, duplications, and splits. The value in the matrix is a $z$-score, calculated as in(Ream, Bankapur, and Friedberg, 2015). As can be seen, a deletion may have happened only in the common ancestor of the $\epsilon$-proteobacteria), few splits (right square) events, and no duplications events in the pairwise comparison of this gene block, showing a high conservation for all event types. Reproduced from (Ream, Bankapur, and Friedberg, 2015) under Creative Common CC-BY-NC 4.0. license. A downloadable higher resolution image is available in `https://github.com/nguyenngochuy91/Ancestral-Blocks-Reconstruction/blob/master/images/Fig4.pdf`

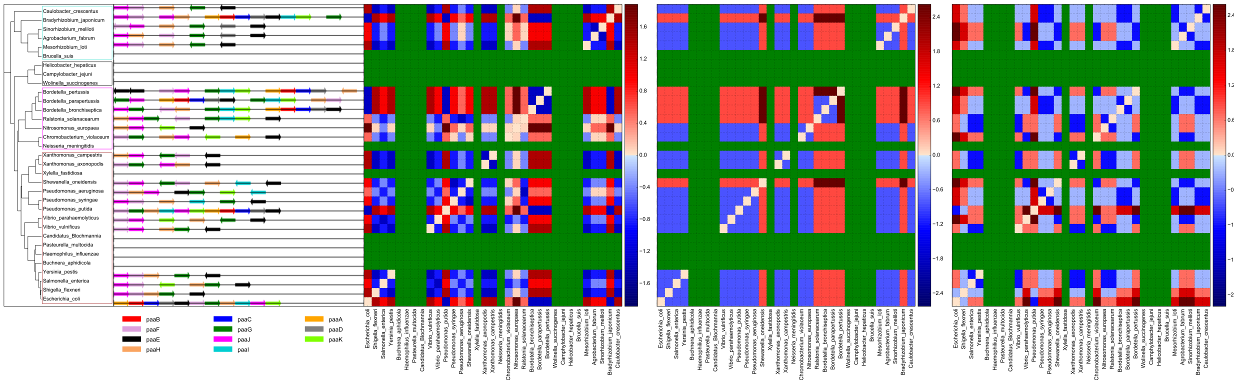


**Fig S4.** Gene block *paaABCDEFGHIJK* phylomatrices, each show the degree of relative conservation of the event between any two species. Left to right: deletions, duplications, splits. Blue to red scale is high-to-low conservation in $z$-score (Ream, Bankapur, and Friedberg, 2015). A downloadable higher resolution figure is available in `https://github.com/nguyenngochuy91/Ancestral-Blocks-Reconstruction/blob/master/images/Fig7.pdf`

#### Gene block *paaABCDEFGHIJK*

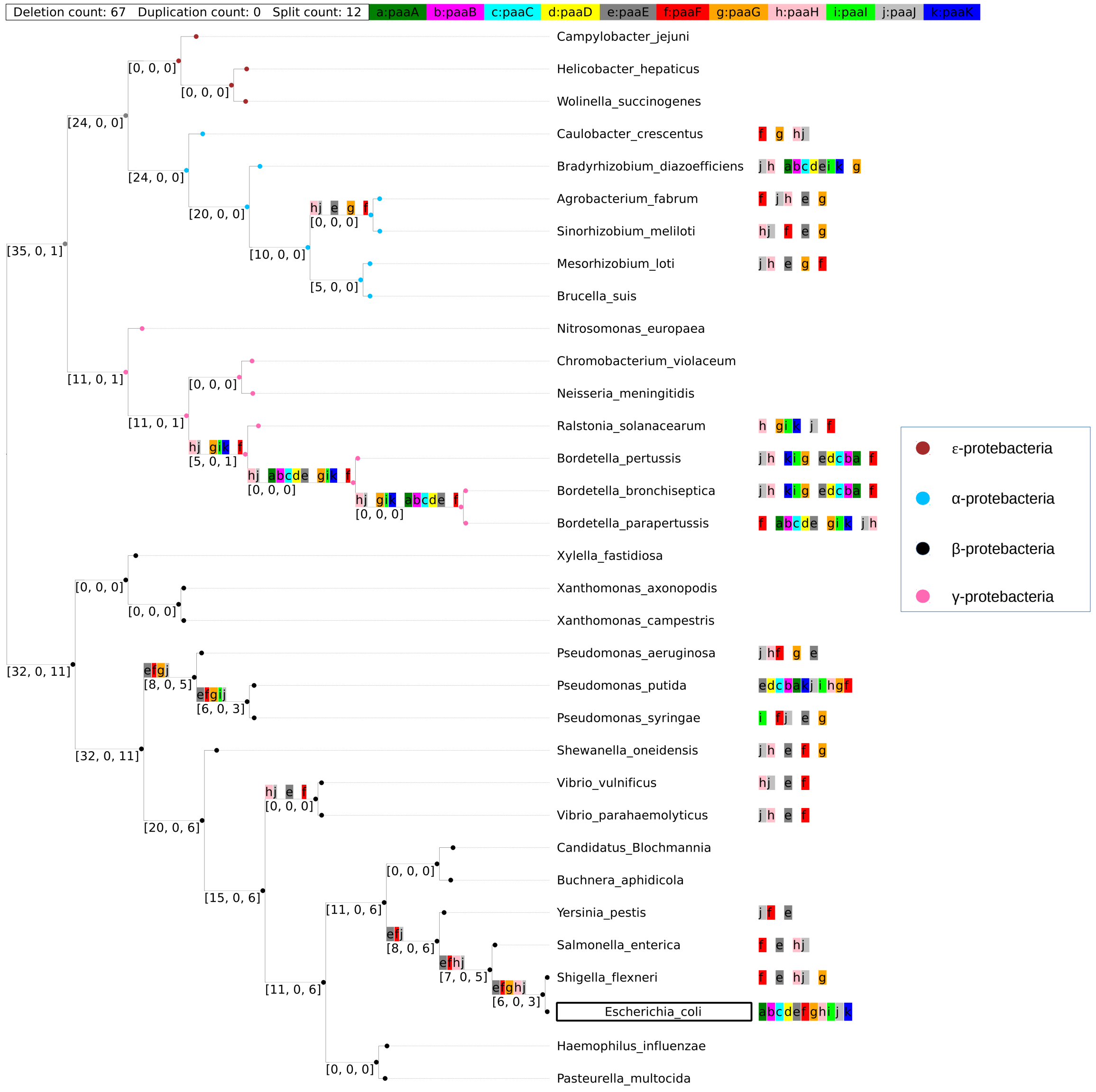


**Fig S5.** Ancestral gene block reconstruction of *paaABCDEFGHIJK* using the local reconstruction approach. Asterisks in front of species names indicate that a minimal orthoblock (which should consist of two or more proximal genes that are orthologs to genes in the reference operon) was not found.
